# A phenotypic brain organoid atlas for neurodevelopmental disorders

**DOI:** 10.1101/2025.09.12.675864

**Authors:** Lu Wang, Yuji Nakamura, Junhao Li, David Sievert, Yang Liu, Toan Nguyen, Prudhvi Sai Jetti, Ethan Thai, Rachel Yibei Zhou, Jiaming Weng, Naomi Meave, Manya Yadavilli, Robyn Howarth, Kevin Camey, Niyati Banka, Charlotte Owusu-Hammond, Chelsea Barrows, Stephen F. Kingsmore, Maha S. Zaki, Eran Mukamel, Joseph G. Gleeson

## Abstract

Thousands of genes are associated with neurodevelopmental disorders (NDDs), yet mechanisms and targeted treatments remain elusive. To fill these gaps, we present a CIRM-initiated NDD biobank of 352 publicly-available genetically-diverse patient-derived iPSCs, along with clinical details, brain imaging and genomic data, representing four major categories of disease: microcephaly (MIC), polymicrogyria (PMG), epilepsy (EPI), and intellectual disability (ID). From 35 representative patients, we studied over 6000 brain organoids for histology and single cell transcriptomics. Compared with an organoid library from ten neurotypicals, patients showed distinct cellular defects linked to underlying clinical disease categories. MIC showed defects in cell survival and excessive TTR+ cells, PMG showed intermediate progenitor cell junction defects, EPI showed excessive astrogliosis, and ID showed excessive generation of TTR+ cells. Our organoid atlas demonstrates both conserved and divergent NDD category-specific phenotypes, bridging genotype and phenotype. This NDD iPSC biobank can support future disease modeling and therapeutic approaches.

**HIGHLIGHTS:**

1. Resource of 352 CIRM-funded genetically-diverse IPSC lines from patients with neurodevelopmental disorders (NDDs).
2. Genome/exome and brain images available for these genetically-diverse IPSC lines.
3. Derived human brain organoids (hBOs) show disease-specific histological and cellular phenotypes.
4. hBO phenotypes show unanticipated differentiation towards non-neuronal cell fates in NDDs.

## INTRODUCTION

Neurodevelopmental disorders (NDDs) occur in more than 4% of children, presenting a range of phenotypes spanning intellectual, motor, language and behavioral deficits, and often including features of autism and epilepsy. NDDs likely reflect defects in cellular, genetic, molecular, and developmental processes, impacting brain architecture encompassing defects in cell fate, migration, survival, and connectivity (Carriba et al., 2023). NDDs often associate with structural brain defects evident on brain imaging, such as reduced cerebral volume evident in microcephaly (MIC), and global excess cortical gyrification, evident in polymicrogyria (PMG). Other NDDs lack clear structural defects, such as non-lesional epilepsy (EPI), characterized by severe recurrent electrographic seizures, or intellectual disability (ID), characterized by cognitive deficits (Spreafico and Tassi, 2012). NDDs present in approximately 25% of all chronic pediatric disorders, collectively imposing an enormous toll, accounting for over $400 billion in annual US health care costs and lost wages (Leigh and Du, 2015).

NDD genetic causes are a topic of intense investigation, with numerous risk factors spanning thousands of genes (Morris-Rosendahl and Crocq, 2020). Specific NDDs can exhibit variable clinical presentations even when caused by the same gene, and conversely, one gene can associate with a range of NDD phenotypes, attributed in part to phenotypic, genetic, and allelic pleiotropy. NDD pleiotropy may be partially explained by cell-type specific gene expression (Gandal et al., 2022; Parikshak et al., 2016). Despite some convergence of early cellular and molecular mechanisms in NDDs, models to capture large-scale genetic and phenotypic diversity remain a distant goal (Velasco et al., 2019; Yoon et al., 2019).

Human brain organoids (hBOs) are three-dimensional induced pluripotent stem cells (iPSCs) derived self-organized cellular clusters, mirroring the structural architectures and cellular complexities of the developing human brain (Kadoshima et al., 2013; Lancaster et al., 2013). Several hBO models of NDDs have uncovered novel cellular and molecular disease mechanisms (Wang et al., 2023b). Shared phenotypes and convergent transcriptomic changes are reported in hBO excitatory neurons in forms of autism spectrum disorders (Jourdon et al., 2023; Paulsen et al., 2022). Although reproducibility of phenotypes is well documented, the robustness of hBOs in modeling disease phenotypes of NDDs across a diverse set of genetic conditions is incomplete.

Starting in 2010s, California Institution of Regenerative Medicine (CIRM) supported the derivation and banking of patient-derived iPSCs from 2184 unique donors across six disease categories and 17 diagnoses, along with matched controls, using best-practice episomal reprogramming (Lin et al., 2020), used to model complex human disease (Dobrindt et al., 2021; Duwaerts et al., 2021; Tegtmeyer et al., 2024). As part of this effort, iPSCs from 352 NDD patients were generated from cell lines we contributed, which passed quality control for genome integrity and pluripotency, and are now available to the scientific community along with whole exome sequencing in most patients. Labs are starting to use these cells in studies ranging from natural variation to viral susceptibility (Wells et al., 2023).

Here we address how well this library represents the spectrum of NDDs, and how well patient hBOs capture disease-specific features. Utilizing this resource, we generated an ‘hBO atlas’ from four major NDD categories, correlating clinical features and genetic results with hBO phenotypes. This atlas consists of over 6,000 hBOs from 10 healthy controls and 35 patients among whom more than half have a clear genetic cause identified. Surprisingly, we found that nearly all patient hBOs showed findings that likely reflect the features of their NDD, with cellular phenotypes tending to cluster based on a broad disease category. Histologic and single cell transcriptomic analysis revealed disease-specific alterations including in non-neuronal cell populations. In particular, we observed a significant increase in astrocyte numbers and evidence of astrocytic reactivation in patients with epilepsy. Additionally, hBOs from patients with microcephaly exhibited excessive TTR (transthyretin) expression and cell survival. These results highlight the robustness of hBO modeling across the NDDs.

## RESULTS

### A comprehensive hBOs phenotypic atlas for NDD

From our cohort of over 10,000 families with likely genetic forms of NDD, we selected 450 from where primary dermal fibroblast cultures were maintained at low passage (P2-P4) from an affected member or members, and at least one parent. Patients were evaluated by a child neurologist and geneticist, brain imaging and biochemical assessments, and assigned to one of four major disease categories: MIC and PMG, representing structural brain defects, and EPI and ID, representing non-structural defects. Patients with MIC had head circumference below two standard deviations for age and brain imaging documenting reduced cerebral volume. Patients with PMG demonstrated global excessive cortical gyrification spaced less than 1.5 cm distance. EPI and ID had minimal to no obvious structural defects on brain imaging. MIC patients showed no PMG, and no PMG patients had MIC, but all patients with MIC, PMG and EPI also displayed some degree of ID, additionally, MIC and PMG patients showed frequent epilepsy, as expected (Spreafico and Tassi, 2012). Patients with ID displayed no documented electrographic seizures. All patients underwent whole genome or exome sequencing to identify likely pathogenic causative mutations. Controls were evaluated by either genome or microarray (See Method and Tables 1 and S1). All relevant clinical and genetic data is available for download (Tables 1 and S1, Zip-File 1, https://brain-org-ndd.cells.ucsc.edu/).

Consented participants had cells were reprogrammed at Cellular Dynamics, Inc., using episomal gene transfer protocol, carried for 10 passages under selection for favorable growth characteristics reflecting pluripotency and expanded (Mack et al., 2011; Takahashi et al., 2007). Of 450 subjects donating samples, 70 fibroblast lines could not be propagated and 28 failed quality control. All remaining lines passed digital karyotyping to exclude most cytogenetic defects, and gene expression array to confirm pluripotency based upon OCT4 and SOX2 expression (See Method, Support File-Figure 1).

**Figure 1.**
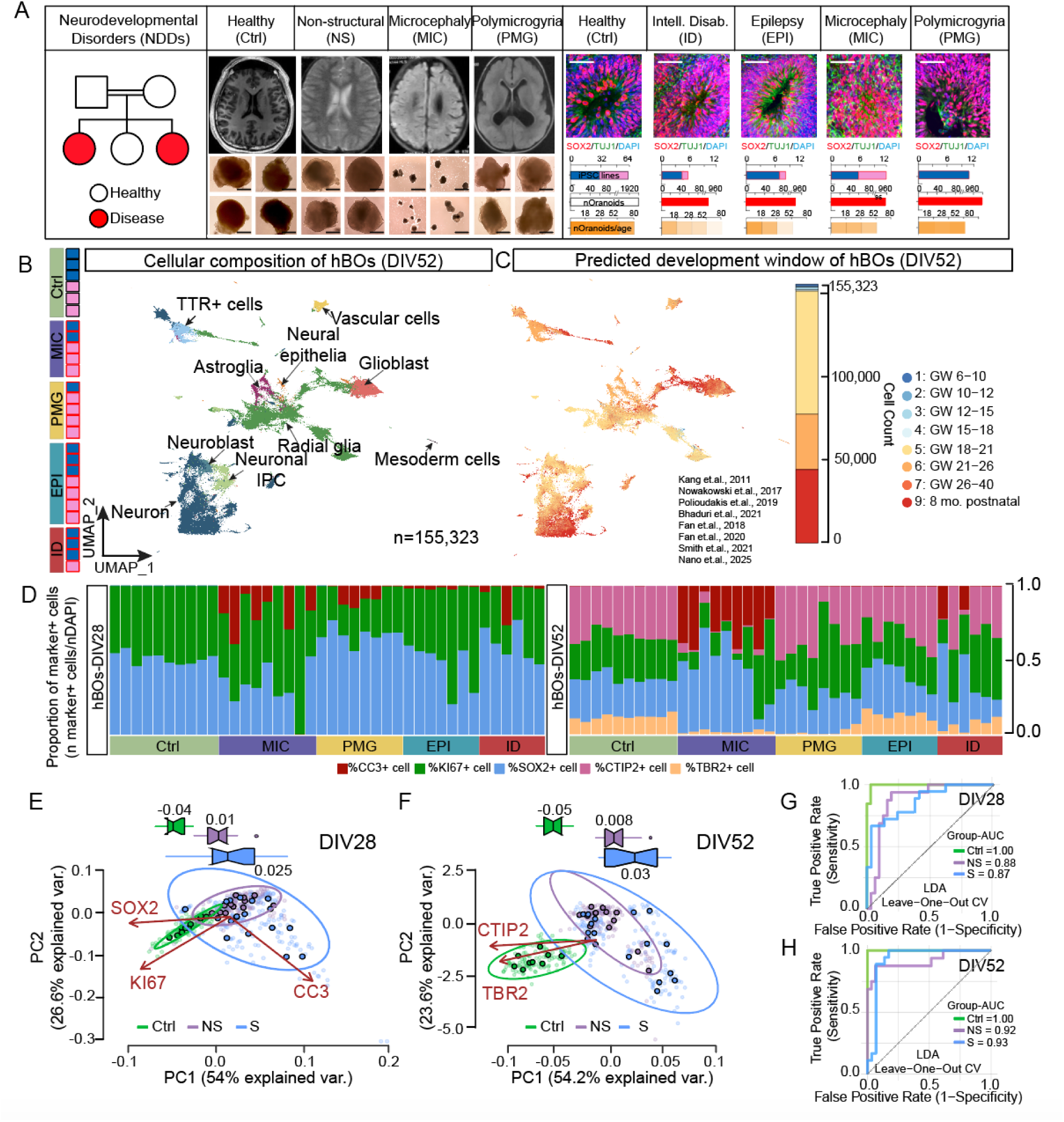
Patient-derived hBOs reveal neurodevelopmental disorder specific cellular phenotypes. (A) Recruitment of families with specific categories of NDD. Consanguinity: double bar. Square: Male; Circle: Female; Red: affected. Subjects include healthy and non-structural brain diseases (Intellectual Disability, Epilepsy) and structural brain diseases (Microcephaly and Polymicrogyria), where brain MRI (top). hBO gross morphology (bottom) correlated with disease class (four representative IPSC lines per class). Bar 0.5mm. Representative rosette histology. Bar 100 μm. Roughly equally represented iPSC lines in males (blue) and females (pink), evaluated at four timepoints, encompassing over 6000 hBOs across all categories, sampled at DIV18, 28, 52, and 80 using immunohistochemistry or single cell sequencing. (B) UMAP of major cell types at DIV52. Left: samples per disease class. Colors: disease categories (i.e. Ctrl: olive, MIC: purple, etc.) used in later figures. Ctrl: black border, Disease: red border. 155,323 cells derived from 26 hBO libraries across five categories assayed. (C) UMAP of hBOs cells mapped onto human fetal brain single cell atlases from gestation week (GW) 6 to 8 months postnatal using label transfer. The hBOs cells predominantly reflect GW18-26 and 8 months postnatal. (D) Barplot of quantitated hBO immunostaining for SOX2, KI67, and CC3 at DIV28 and DIV52, and CTIP2 and TBR2 at DIV52. Percentage of marker-positive cells calculated for each replicate (technical and biological). Mean calculated per individual iPSC line, using DAPI+ normalization. Raw data in Table S2. (E-F) PCA summary of hBOs immunostaining using SOX2, KI67, and CC3 at DIV28 (E) and CTIP2 and TBR2 at DIV52 (F). Mean contribution of variance for PC1 for each category (Ctrl, nonstructural (NS), and structural (S). Percent of variance explained along *x* and *y* axes. Twelve biological replicates and three image regions per section quantitated. Replicate: small dots. Replicates per iPSC line: big dots. Raw data in Table S2. (G-H) Performances of linear discriminant analysis (LDA) in classifying samples from diagnostic categories (Ctrl, NS, and S) for DIV28 (G) and DV52 (H). LDA classifiers were trained with leave-one-out cross-validation using the arcsine-transformed proportion of cells expressing markers as a percent of DAPI+ nuclei per imaging field at DIV28 (CC3, KI67, and SOX2) and DIV52 (TBR2, CC3, KI67, SOX2, and CTIP2). Multi-class Receiver Operating Characteristic (ROC) curves of the LDA classifiers for DIV28 (G) and DIV52 (H). AUC, area under the curve.

We selected 35 patients, representing the four disease categories (9 from MIC, 8 lines from PMG, 7 lines from EPI, 11 lines from ID), and 9 control lines from unaffected parents. We also included the H1-human embryonic stem cell line, to allow comparison with prior studies (Pollen et al., 2019). Genomes were reviewed by qualified specialists, revealing a likely genetic cause in 24 patients (68.5%) (Tables 1 and S1). To confirm pluripotency of this library, iPSC lines used for hBO generation, as well as an additional random 14 lines were validated for OCT4 staining (Support File-Figure 1). We next generated over 6000 hBOs using a single widely adopted method, and harvested hBOs at days in vitro (DIV) 28 and 52 for histological analysis for all lines, and DIV52 for single cell transcriptomic analysis in approximately half of the lines (Figure 1A)(Kadoshima *et al*., 2013).

### hBO as a robust model for early brain development

We first characterized hBOs variability from the ten healthy iPSC lines (H1 and 9 healthy parent lines), matched for sex (Tables 1 and S1). We measured diameter of ten percent of all hBOs and found minimal variability at DIV28 and 52 (2.1 +/- 0.2 mm vs. 3.9 +/- 0.4 mm, respectively), in keeping with recent publications (Table S2)(Bhaduri et al., 2020; Quadrato et al., 2017).

At DIV28 we found nearly identical expression patterns and spatial locations for radial glial cells (RGCs) and immature neurons, evidenced by SOX2 and TUJ1 immunostaining, respectively (Figures S1A1-10). Approximately 56.57% nuclei per section were KI67+, and 0.03% were CC3+ at DIV28, consistent with rapid cell expansion (Figures S1B1-10), and apical surfaces evidenced ZO1 localization, denoting apical junctions (Figures S1C1-10). At DIV52 neural rosettes evidenced flower-like appearance laminated with SOX2+ progenitors surrounded by CTIP2+ deep cortical layer neurons and TBR2+ intermediate progenitor cells (Figures S1D1-10). Quantifications for each of cellular feature assayed at DIV28 and 52 showed limited variation (Figures S1E-F, Table S2, See Method). We extended some cultures to DIV80 but observed more variability in distribution of TBR1+ cells and a dissolution of laminated architecture, and for this reason we limit most analysis to earlier timepoints (Support File-Figure 2).

**Figure 2.**
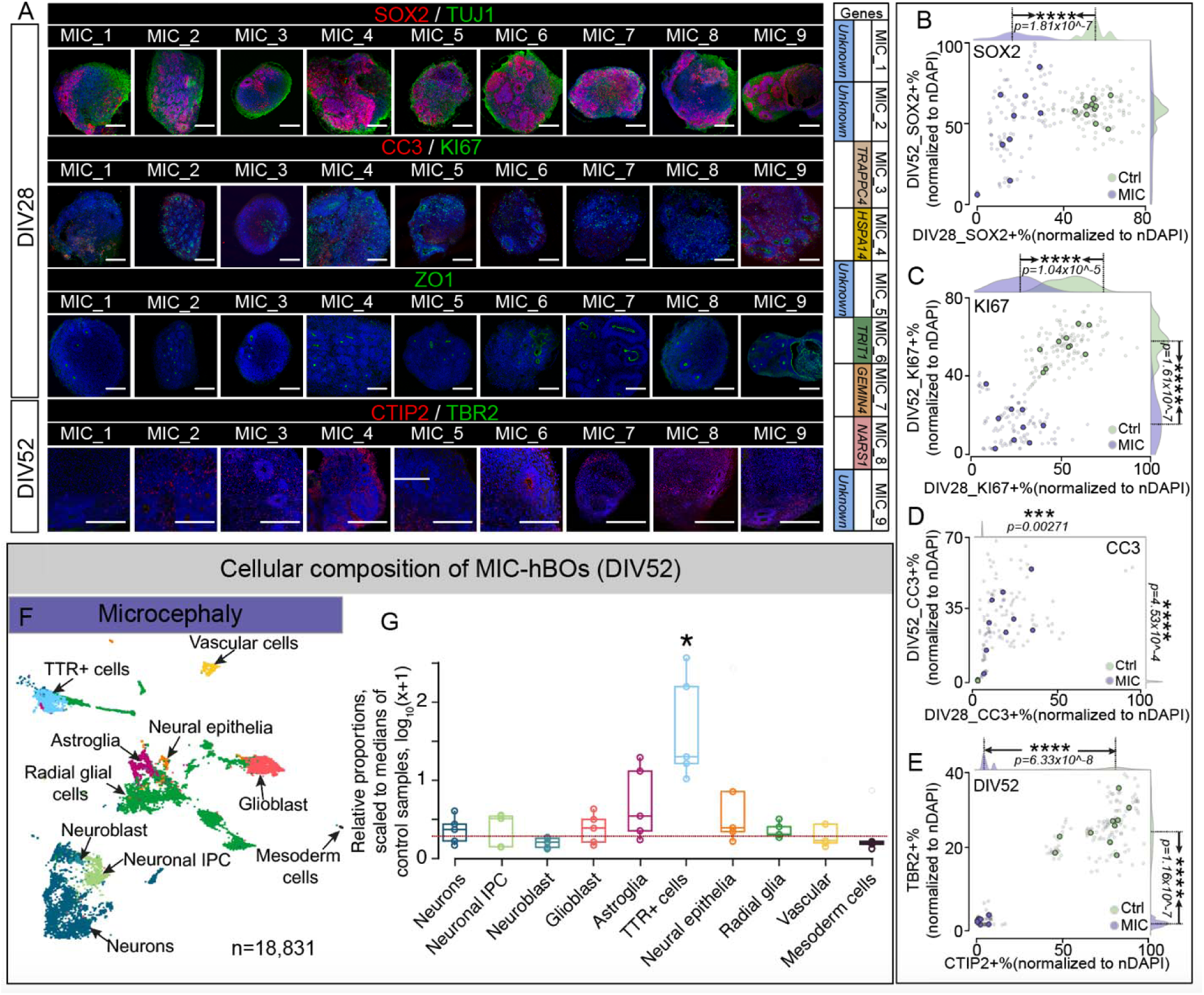
hBOs of patients with microcephaly exhibit fate shifted toward TTR+ cells. (A) Similarities in 9 MIC patients at DIV28 and DIV52, four with established genetic cause (noted at right, e.g., *TRAPPC4*). In comparison with 10 Ctrl (Fig. S1), MIC showed altered neural rosette composition. SOX2 (radial glial progenitors), TUJ1 (immature neurons), CC3 (apoptosis), KI67 (proliferation), ZO1 (apical tight junctions), CTIP2 (deeper layer neurons), TBR2 (intermediate progenitors), DAPI (nuclear, for normalization). Bar: 400μm. (B-E) Reduced SOX2+ cells at DIV28 (*x-*axis) with less noticeable differences at DIV52 (*y-*axis) in (B). KI67 was significantly decreased at both timepoints in (C). (D) Few if any CC3+ cells observed in Ctrl (all dots at bottom left) whereas patients showed significant increase, especially at DIV52. (E) Few if any CTIP2+ or TBR2+ cells observed at either timepoint in all patients. Two-sided t-test, *p<0.1, **p<0.01, ***p<0.001, and ****p<0.0001. Smaller dots: percentage of DAPI positive cells for each organoid (ten control lines and nine MIC lines, twelve organoids derived from each line as technique replicates). Larger dots: mean of the percentage-positive cells from all organoids derived from the same iPSC line. *p* values: differences among individual iPSC lines (See detailed p-values in Table S7). (F) UMAP cellular clusters in MIC at DIV52 from 18,831 cells in MIC. (G) Cellular fractions for each cluster showed significant increase in TTR+ cells in MIC. *, FDR = 0.024. Horizontal dashed line (red): normalized value FDR (log_10_2) where there were no differences between the two groups.

We used scRNA-seq to profile hBOs from approximately half of the controls and patients, generated from 26 individuals (6 controls and 20 patients), assessing 155,323 cells. We documented 59 cellular clusters, annotated into 10 major classes, corresponding to reported major human fetal brain cell types (Figures 1B and S2A-D, Tables S3-5, Support File-Figures 3-4). Standard label transfer methods showed hBOs cells corresponded to brain developmental stages spanning gestation week (GW) 6 through 8 months postnatal, with a enrichment in GW18-26 at the second trimester and 8 months postnatal, highlighting neurogenesis (Figures 1C and S3A, See Method) (Bhaduri et al., 2021; Fan et al., 2018; Fan et al., 2020; Nano et al., 2025; Nowakowski et al., 2017; Polioudakis et al., 2019; Smith et al., 2021; Zhong et al., 2018). We found features reflecting several anatomical brain regions, predominantly forebrain, but also diencephalon, as noted by others (Figure S3B, Table S3, See Method)(Braun et al., 2023; Kadoshima *et al*., 2013). These results suggest that hBOs from both controls and patients capture major steps in neurogenesis and maturation of developing human brain in a spatial and temporal specific manner.

**Figure 3.**
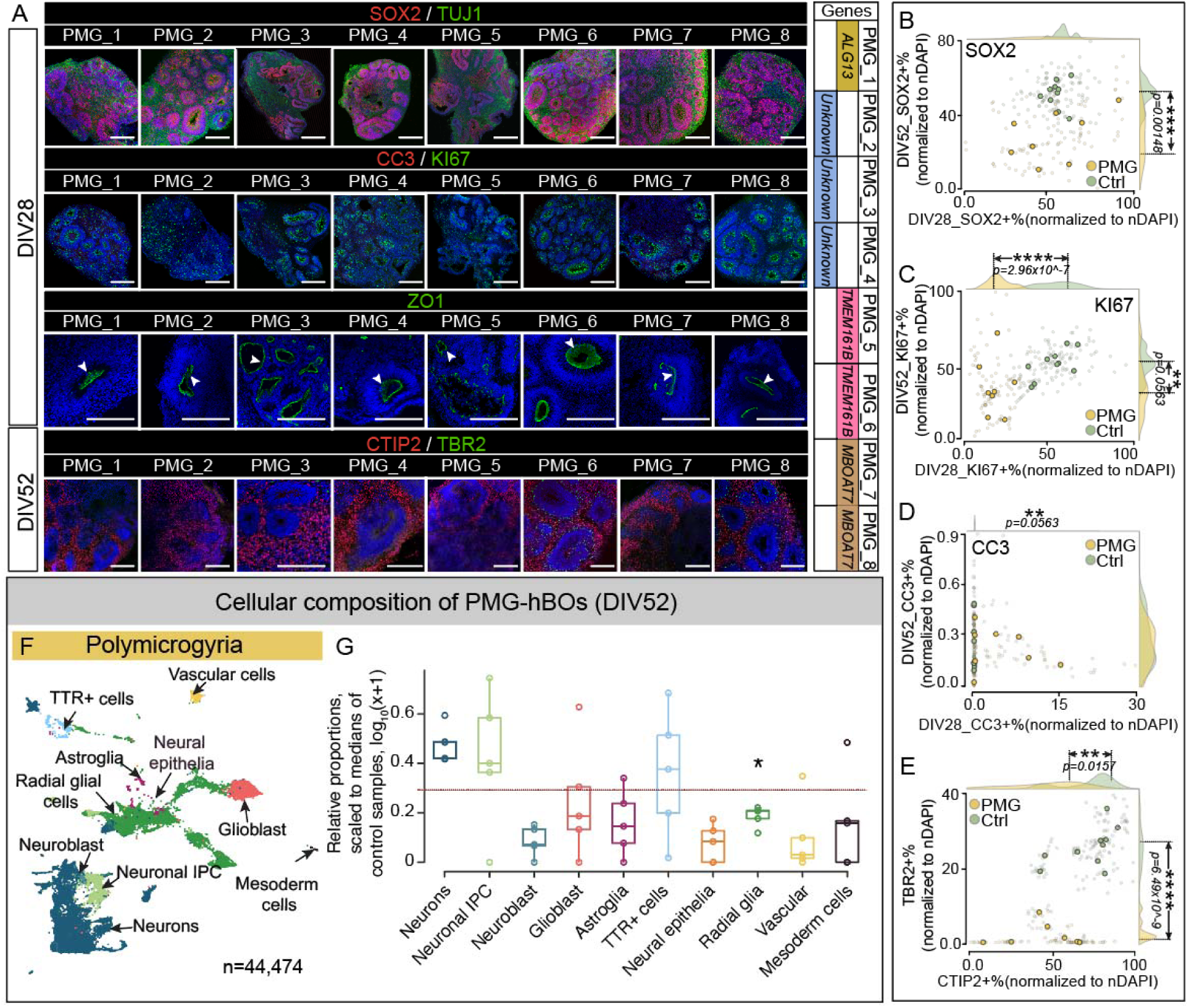
hBOs of patients with polymicrogyria exhibit defects of intermediate progenitor cell-cell junctions. (A) hBOs derived from 8 PMG patients at DIV28 and DIV52, six with established genetic cause (noted at right, e.g., *TMEM161B*), in comparison with 10 Ctrl (Figure S1). Bar: 400um. Bar plot on left: pink: Female; blue: Male. (B-E) Mildly reduced SOX2+ cells at DIV28 (*x-*axis) with less noticeable differences at DIV52 (*y-* axis) in (B). KI67 significantly decreased at both timepoints in (C). (D) Few if any CC3+ cells in Ctrl (all dots at bottom left) whereas PMG showed significant increase, especially at DIV52. (E) marked reduction in CTIP2+ and TBR2+ cells at either timepoint in all patients. Two-sided t-test, *p<0.1, **p<0.01, ***p<0.001, and ****p<0.0001. Smaller dots: percentage of DAPI positive cells for each organoid (ten individual control lines and eight individual PMG lines, twelve organoids derived from the same iPSC line as technical replicates). Larger dots: mean percentage-positive cells from all hBOs derived from the same iPSC line. *p* values: differences among individual iPSC lines (See detailed p-values in Table S7). (F) UMAPs: Cellular clusters from 44,474 cells from PMG at DIV52. (G) Cellular fractions for each cluster in both categories showed subtle non-significant differences (*), FDR = 0.178. Red dashed line: normalized value FDR (log_10_2). No differences between the two groups.

### hBOs reproducibly mirror cellular phenotypes of NDDs

To capture cellular NDD phenotypes, we immunostained hBO sections from patients to quantify numbers of cells expressing SOX2, CTIP2, TBR2, CC3 and KI67 as a percent of DAPI+ nuclei per imaging field at DIV28 and DIV52 (Figure 1D and S4A, Table S2, See Method). We aggregated hBOs findings from patients with and without structural brain defects, using principal component analysis. At DIV28, patients and controls differed significantly (evident by explained PC1 at -0.04 for controls, 0.025 and 0.01 for patients with or without structural brain defects), showing limited overlapping distributions of immunopositive cells. However, patients as a whole partially or fully overlapped with each other at this early timepoint (Figure 1E, Table S2). At DIV52, hBOs from patients with structural brain defects showed greater separation from those without (Figure 1F, Table S2, explained PC1 at 0.03 and 0.008 from patients with or without structural brain defects). We next leveraged a linear discriminant analysis to classify the samples with ‘leave-one-out’ cross-validation. This analysis allowed classification of disease categories based upon cellular phenotypes, with a robust discriminatory performance as indicated by area under the curve (AUC) of 0.88 and 0.87 for DIV28 and 0.92 and 0.93 for DIV52 for patients with and without structural brain defects (Figures 1G-H and S4B, Tables S2 and S6). However, the principal component analysis underperformed at DIV80 (Support File-Figure 2B) based on explained PCs, likely reflecting the greater variability evident at later timepoints. These findings suggest hBOs capture aspects of diverse NDDs at cellular levels, that patient-hBOs can be most often distinguished from controls, and can most often distinguish patients with and without structural brain defects, depending upon the timepoint of evaluation.

### Shift in fate from neuron to TTR+ cells in hBOs from patients with microcephaly

Focusing initially on MIC, we studied hBOs from nine patients, referred to as MIC-hBOs, of which five showed biallelic likely disease-causing mutations. MIC_3 was mutated for *TRAPPC4*, MIC_6 was mutated for *TRIT1*, MIC_7 was mutated for *GEMIN4*, MIC_8 was mutated for *NARS1* (Aldhalaan et al., 2021; Ghosh et al., 2021; Kernohan et al., 2017; Van Bergen et al., 2020; Wang et al., 2020). The remaining four lines had unknown genetic causes although MIC_4 had a biallelic damaging mutation in the candidate gene *HSPA14*, not yet annotated in OMIM (Tables 1 and S1).

hBO diameter in microcephaly was grossly reduced compared with controls, first apparent at DIV18, supporting prior observations (Figure S5A, Table S2)(Lancaster *et al*., 2013). These differences became even more apparent at DIV52 (1.9 +/- 0.29mm vs. 3.9 +/- 0.4 mm at DIV52, *p*=2.18x10^-38), suggesting impaired hBO growth likely reflecting defects in cell survival or proliferation (Carpentieri et al., 2022)(Figure S5B, Table S2).

We found that both SOX2+ and KI67+ cells, normally tightly clustered around rosettes, were instead scattered across the entire hBO (Figures 2A and S1A-B). ZO1, normally present at apical surfaces, was dispersed and at reduced levels, and very few lines showed usual populations of CTIP2+ or TBR2+ cells (Figures 2A and S1C-D). Plotting percent SOX2+ cells in microcephaly vs. control at DIV28 showed good stratification, whereas at DIV52, SOX2+ showed broader but largely overlapping distributions (*p*=1.81x10^-7 at DIV28, Figure 2B, Table S7). Percent KI67+ cells distinguished microcephaly vs. control at both timepoints (DIV28 *p*=1.04x10^-5, DIV52 *p*=1.61 x10^-7, Figure 2C). Percent CC3+ cells were much greater in microcephaly vs. control at both DIV28 (*p*=0.00271) and DIV52 (*p*=4.53x10^-4), suggesting reduced cell survival (Figure 2D). Percent CTIP2+ and TBR2+ cells were both dramatically reduced at DIV52 (CTIP2+ *p*=6.33x10^-8, TBR2+ *p*=1.16x10^-7, Figure 2E), suggesting failure to accumulate intermediate progenitor cells and deep-layer cortical neurons, consistent with defects in both cell survival and proliferation.

Since hBOs in microcephaly represented at least 5 different genetic causes, we selected one line carrying each mutation for scRNA-seq with these hBOs at DIV52. UMAP analysis of 18,831 cells in microcephaly patients showed 10 major cell types (Figure 2F), but neural cell types (i.e., neurons, neuroblast, and neuronal intermediate progenitors) in microcephaly were reduced approximately 74% (cell type proportion in scRNA-seq), consistent with our immunostaining results (Figure 2A, Tables S3 and S7). This reduction appeared to be driven in part by a reciprocal emergence of a ‘TTR+’ cell population (Figures 2G and S5C, FDR=0.024, Table S8). This data suggests depletion of specific neural classes and concurrent expansion of TTR+ cell types in microcephaly.

We assessed the gene expression for the five cellular marker genes in each of the cell subclasses observed from single cell RNA-seq by gene set enrichment analysis. We observed several pathways associated with functions for mitochondrial and ATP metabolism, which were positively enriched in glioblast and VGlut2+ inhibitory neurons, while negatively enriched in TTR+ cells and radial glial cells (Figure S5D, Tables S9-10, See Method).

In addition, we assessed the expression of genes that encode the protein markers we measured with immunostaining in each of the cell clusters and observed decreased *MKI67* (encoding Ki67, targeted Wilcoxon tests, two-sided raw p-value = 0.0403, Table S11) in neuroblast, consistent with loss of proliferation (Figure S5E)(Qian et al., 2016).

### Defects in cellular adhesion in neuronal IPCs in hBOs from patients with polymicrogyria

With the same strategy, we characterized hBOs derived from eight individuals with polymicrogyria. PMG_1 was mutated for *ALG13*, PMG_5 and PMG_6 shared mutations in *TMEM161B*, PMG_7 and PMG_8 shared mutations in *MBOAT7* (Akula et al., 2023; Johansen et al., 2016; Wang et al., 2023a), while the remaining three lines had unknown causes (Tables 1 and S1).

Unlike microcephaly, hBOs from patients with polymicrogyria showed grossly normal size, but revealed consistent defects in apical junctions and intermediate progenitor numbers (Figure 3A, Table S2). We noted similar percent SOX2+ cells at DIV28, but significant reduction at later timepoint DIV52 (*p*=0.00148, Figures 3A-B). KI67 staining was localized near the apical surface like Ctrl but was reduced in percent at both timepoints (DIV28 *p*=2.96x10^-7, DIV52 *p*=0.0563, Figures 3A and C, Tables S2 and S7). ZO1, which typically labels apical junctions marking concentric ventricular-like zones, were instead often discontinuous, fragmented, and compressed (Cakir et al., 2019) (Figures 3A and S1C). While percent CC3+ cells were slightly increased, TBR2+ cells showed a severe reduction (*p*=6.49x10^-9, Figures 3D-E). These data reveal depletion of intermediate progenitors, and apical junction defects.

Since these hBOs derived from patients with at least four different genetic mutations, we selected lines carrying each mutation for scRNA-seq with these hBOs at DIV52. UMAP analysis of 44,474 cells identified similar clustering of the 10 major cell type groupings, with only subtle differences in cellular populations (Figures 3F-G and Table S8). Because there were no major differences in cell types, we performed subclustering of neuronal intermediate progenitors, which, from TBR2 staining, was reduced (Figure S6A). We found the neuronal intermediate progenitor cluster was expanded in proportion, in part likely due to a significant increase in DLX2+ cells (Figures S6A-B and Table S8, *p*=0.08), supporting disruption in neuronal intermediate progenitor accumulation. Moreover, pseudobulk analysis followed by gene set enrichment analysis suggested several pathways associated with ribonucleoproteins and mitochondrial stress, which were positively enriched in TTR+ cells, glioblast and vGlut-2+ inhibitory neurons, while negatively enriched in radial glial cells (Figure S6C, Table S9-10).

### Enrichment of astrocytes in hBOs from patients with epilepsy

We next characterized hBOs derived from seven individuals with severe epilepsy. Among these, EPI_2 was mutated for *MDGA2*, EPI_4 was mutated for *FAM57A*, EPI_5 was mutated for *ZBTB40*, while the remaining four lines had unknown genetic causes(Connor et al., 2016; McCammon et al., 2017; Ward et al., 2024)(Tables 1 and S1).

hBOs from patients with epilepsy were grossly within normal limits but were slightly smaller in size (*p*=4.57X10^-5 at DIV28, *p*=0.000443 at DIV52, Table S2). Immunostaining revealed significant reductions in percent of SOX2+ and KI67+ cells at DIV28 (*p*=0.00102 and *p*=9.07x10^-5, respectively, Figures 4A-C, Table S7). CC3+ cells were increased significantly at DIV28 (*p*=0.0122), while no significant differences were detected for CC3+ cells at DIV52 (Figure 4D). there were notable decreases in percent of both CTIP2+ and TBR2+ cells at DIV52 (*p*=6.7x10^-6 and *p*=1.26x10^-5, respectively) but not at DIV28 (Figure 4E), suggesting a delay in neuronal maturation.

**Figure 4.**
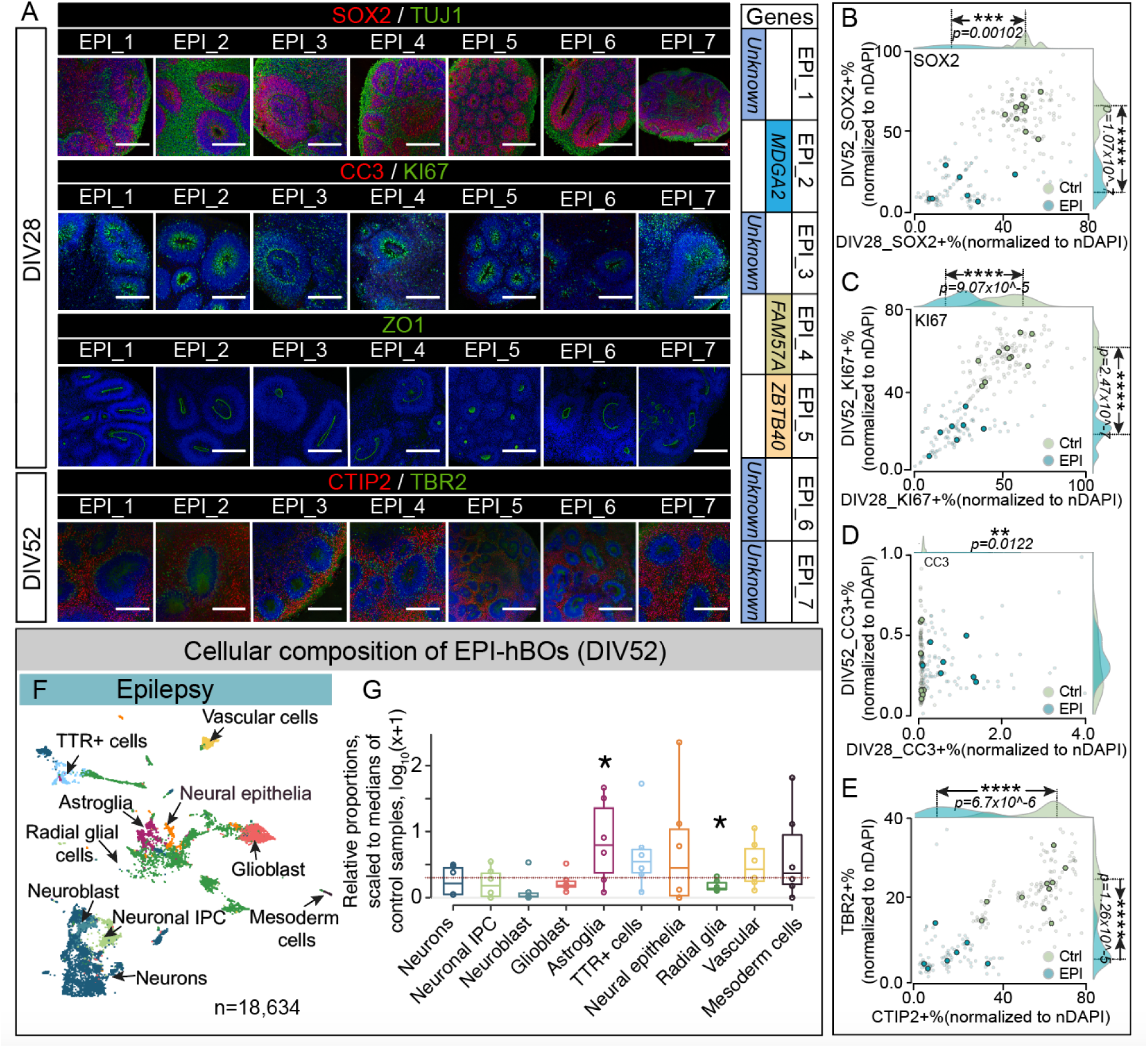
hBOs of patients with epilepsy exhibit enrichment of astroglia cell proportions. (A) hBOs from seven patients at DIV28 and DIV52, five with established genetic cause (noted at right, e.g., *ZBTB40*). Comparison with 10 Ctrl (Figure S1) showed differences in neural rosette composition at DIV52 with greater CC3+ and fewer CTIP2+ and TBR2+ cells. Bar: 400um. Bar plot on left: pink: Female; blue: Male. (B-E) Scatter plots: reduced SOX2+ cells at DIV28 (*x-*axis) with less noticeable differences at DIV52 (*y-*axis) (B). KI67 significantly decreased at both timepoints (C). (D) Few if any CC3+ cells in Ctrl (all dots at bottom left) whereas patients show significant increase, especially at DIV52. Two-sided t-test, *p<0.1, **p<0.01, ***p<0.001, and ****p<0.0001. Smaller dots: percentage of DAPI positive cells from the iPSC lines assayed (ten individual control lines and seven individual EPI lines, twelve organoids derived from the same iPSC line as technique replicates). Larger dots: mean positive cells from each organoid derived from the same iPSC line. *p* values: differences among individual iPSC lines (See detailed p-values in Table S7). (F) UMAPs: Cellular clusters from 18,634 cells from EPI at DIV52. (G) Cellular fractions for each cluster revealed statistical increases in astroglia (cherry). *, FDR = 0.087 for astroglia and radial glia. Horizontal red dashed line: normalized value FDR (log_10_2) where there were no differences between the two groups.

Since hBOs from patients with epilepsy represented only 3 established genetic causes, we performed scRNA-seq with DIV52 hBOs derived from all lines to ensure representation across a diversity of patients. UMAP analysis of 18,634 cells showed similar clustering of 10 major cell groupings but revealed significantly increased astroglia (Figures 4F-G, Table S8). These data were confirmed using immunostaining of GFAP+ cells in hBOs, where we observed an activated astrocytic morphology and significant increase in GFAP+ cells compared to control (Figures S7A-B, Table S2). Pseudobulk analysis followed by gene set enrichment suggested altered pathways and gene expression across different cell types, including significant astroglial upregulation of *EMP1* reported to regulate astroglia functions (Figures S6D and S7A-E, Tables S9-10)(Liddelow et al., 2017)

Astrogliosis has been reported in histopathological brain sections in patients with epilepsy, obtained either postmortem or after lesional resection surgery (Vezzani et al., 2022). To confirm that the astrogliosis we observed in hBOs is clinically relevant, we performed immunohistochemistry for GFAP in human neocortical sections obtained from lesional epilepsy resections. From two patients evaluated, we observed striking accumulation of GFAP+ cells scattered diffusely throughout the section, enriched in perivascular regions (Figures S7F-G, Table S2). These results support the clinical relevance of astroglia accumulation in hBOs from patients with epilepsy.

### Neurotransmitter system expression defects in TTR+ cells in hBOs from patients with intellectual disability

Finally, we characterized hBOs derived from six individuals with intellectual disability, termed ID- hBOs. Among these, ID_1 was mutated for SLC4A10, ID_2 was mutated for FAT1, ID_3 was mutated for *ALDH18A1*, ID_4 was mutated for *CDH20*, ID_5 and ID_6 was mutated for *MAST3* (Tables 1 and S1)(Driessens et al., 2023; Fasham et al., 2023; Shu et al., 2021).

In contrast to other categories, intellectual disability hBOs exhibited considerable variability in histological phenotypes, but within each patient line was highly reproducible. On average, hBOs were significantly smaller and showed overall decrease in neural cell populations, although not as severe as in microcephaly (Table S2). The percent of SOX2+ and KI67+ cells were consistently reduced in all ID lines (Figures 5A-C and S1A-B), whereas percent of CC3+ cells was increased at both DIV28 and DIV52 (Figures 5A and C-D). ZO1 displayed either a clear concentric apical location, as in ID_1, 5 and 6, or was fragmented or compressed as in ID_2, 3 and 4 (Figures 5A and S1C), likely reflecting a diversity of mechanisms in early neural events in ID. While CTIP2+ and TBR2+ cells were localized correctly, the percent of cells of each cell type was reduced, suggesting potential delayed differentiation (*p*=0.0013 and *p*=9.79x10^-6, respectively, Figures 5A, E and S1D, Tables S2 and S7). Results from five additional ID patients, mutated for different genes (ID_8 mutated for *FN3K,* ID_9 mutated for *HIATL1,* ID_10 mutated for *SPATC1*, and ID_11 mutated for *ATP8A2)* were similar (Figure S8A-D, Tables 1 and S1-2).

**Figure 5.**
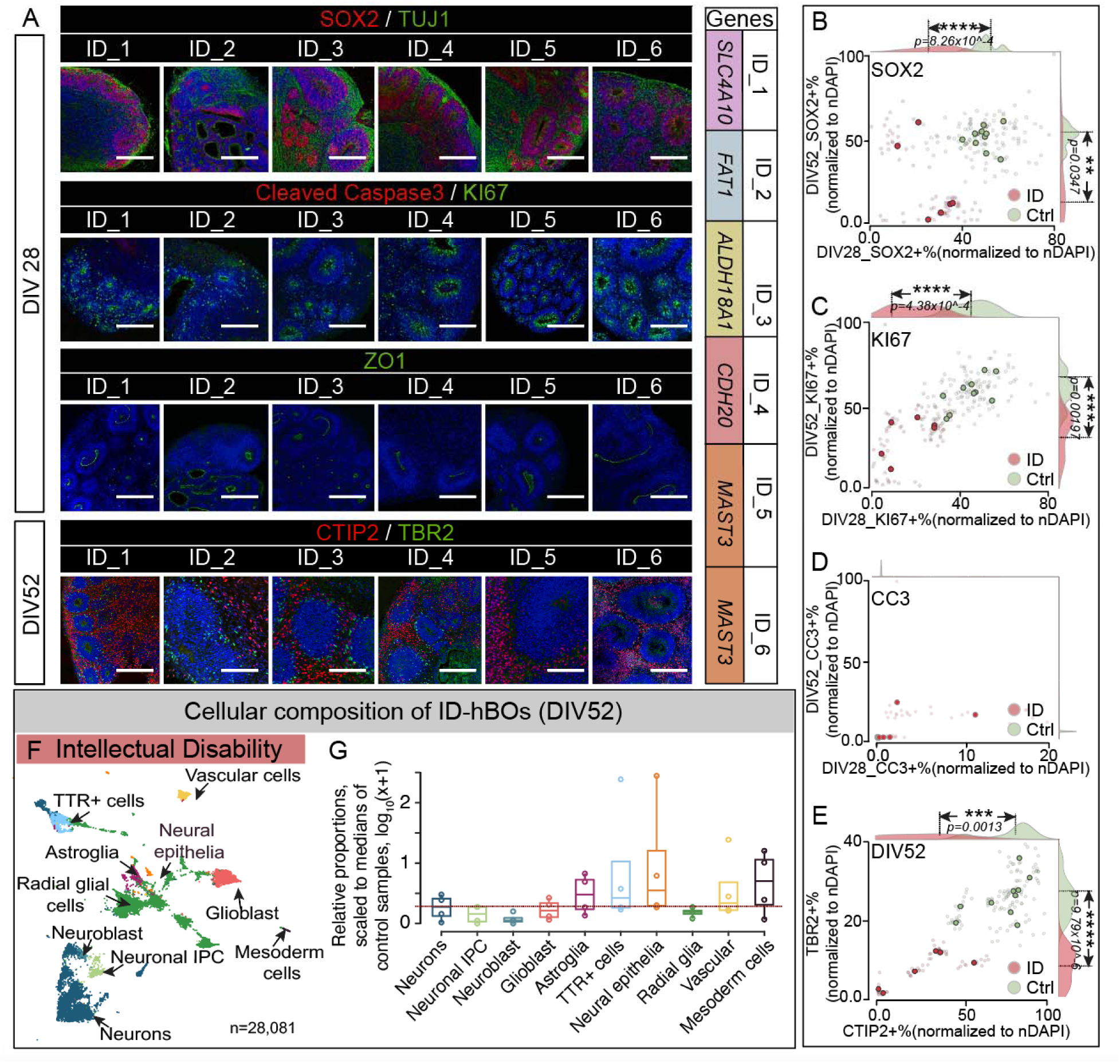
hBOs of patients with intellectual disability exhibit excessive TTR+ cells. (A) hBOs derived from six patients with intellectual disability at DIV28 and DIV52, five with established genetic cause (noted at right, e.g., *ALDH18A1*). Comparison with 10 Ctrl (Fig. S1) showed apparent differences in neural rosette composition at DIV 52, with fewer CTIP2+ and TBR2+ at DIV52. Bar: 400um. Bar plot on left: pink: Female; blue: Male. (B-E) Significantly reduced SOX2+ cells at DIV52 (*x-*axis) in (B). KI67 showed significant decrease at both timepoints in (C). (D) increased CC3+ cells in EPI vs. Ctrl at both timepoints. (E) significant reduction in CTIP2+ and TBR2+ cells at DIV52. Two-sided t-test, *p<0.1, **p<0.01, ***p<0.001, and ****p<0.0001. Smaller dots: percentage of DAPI positive cells for each hBO (ten control lines and six ID lines. Twelve organoids derived from the same iPSC line as technique replicates). Larger dots: mean percentage-positive cells from all hBOs derived from the same iPSC line. *p* values: differences among individual iPSC lines (See detailed p-values in Table S7). (F) UMAP clusters in ID hBOs at DIV52. Total often clusters captured (28,081 cells in ID-hBOs analyzed). (G) Significant increase in TTR+ cells in ID. Fractions of each cell class for each ID individual, normalized to the median of proportion across Ctrl individuals. Horizontal red dashed line: normalized FDR (log_10_2). No differences between the two groups.

scRNA-seq analysis across 28,081 cells at DIV52 carrying various gene mutations of intellectual disability patients, profiled by UMAP, revealed a slight increase in percent of TTR+ cells (Figures 5F-G, Table S7). Pseudobulk followed by gene set enrichment analysis suggested altered pathways, including enrichment of transforming growth factor beta in VGlut2+ neurons (Figure S6E and Tables S9-10).

### CRISPR mediated HDR correction of patient mutations rescues hBO phenotypes

We wondered whether the pathogenic mutations were solely responsible for the hBO phenotypes. We thus selected iPSCs from two patients, i.e. microcephaly (due to *NARS1* mutation, c50C>T) and intellectual disability (due to *ACTL6B* mutation, c.460C>T), and performed CRISPR correction using homology directed repair. A silent mutation (c.48C>T for *NARS1* correction and c.465C>T for *ACTL6B* correction) was introduced to disrupt the PAM sequence and confirm zygosity (Simkin et al., 2022)(Figure S9A). Multiple clones of the corrected lines were propagated, and confirmed for genetic correction by Sanger sequencing, and for pluripotency by OCT4 staining (Figures S9B-C and S10A-B, Table S12).

We generated hBOs from five corrected clones for *NARS1* mutations; three of these clones were homozygous (clone 90, 65, and 37), and two were heterozygous corrected (clone 27 and 53), but since carriers for these mutations were healthy, we expected both corrections should rescue phenotypes. We found all hBOs with corrected *NARS1* exhibited sizes comparable to neurotypicals (Figures S9D-E, Tables S2 and S12). We next generated hBOs from two corrected clones for *ACTL6B,* both homozygous corrected. *ACTL6B* mutant hBOs show expression abnormalities of the early response genes: *FOS*, *FOSB*, *DUSP5*, *VGF*, *FOSL2*, and *JUN* (Wenderski et al., 2020)(Figure S10C, Tables S2 and S12). CRISPR correction showed rescue of these early response gene misexpressions. These data are consistent with rescue of phenotypes with CRISPR correction.

## DISCUSSION

Here we present an inexhaustible patient-derived iPSC resource representing major structural and non-structural forms of NDDs, part of a larger CIRM-funded iPSC library. This genetically diverse cohort was recruited from 8 different countries, and is coupled with detailed clinical and radiographic images, as well as WGS or WES on each participant (Table S1 and Zip File 1, See Method). Analysis of these lines uncovered both disease- and mutation-specific cellular and transcriptional phenotypes that could be corrected by CRISPR editing. The resource, available to the research community, also includes lines from unaffected healthy family members to serve as genetically similar controls, capturing populational level genetic diversity and genetic background in NDDs.

Because NDDs can show both overlapping and disease-specific clinical presentations, we divided the cohort into major clinically-relevant subtypes: structural vs. nonstructural brain disease, and then further divided into four categories. While underlying gene-specific mechanisms will require more focused future efforts, we identified phenotypes that likely represent developmental defects across NDDs. For instance, we found expansions in TTR+ cells in microcephaly and intellectual disability categories and astrogliosis in epilepsy. These findings were further studied in single cell transcriptome analysis, uncovering both convergent and divergent cellular phenotypes.

### Emergence of disease-specific cellular phenotypes

Our results highlight the potential for hBOs to capture cellular phenotypes across a range of heterogeneous monogenetic NDDs. This effort was supported in part by a greater uniformity of hBO generation from standardization in protocols and reagents used here (Velasco *et al*., 2019; Yoon *et al*., 2019). Nearly all patient lines analyzed showed spatial and temporal cellular differences that distinguished them from controls, which suggest a greater potential to model disease and uncover basic mechanisms. However, NDDs are clearly genetically complex, subject to gene-gene interactions, gene-environmental interactions and stochastic factors, only a subset of which could be adequately modeled here.

### Microcephaly

Patients with microcephaly show reduced size of head and brain (Jayaraman et al., 2018). Many studies document impaired cell division, proliferation, and excessive death in both hBOs and mouse models (Chai et al., 2021b; Dhaliwal et al., 2021; Fair et al., 2022; Lancaster et al., 2013; Urresti et al., 2021; Wang et al., 2020; Zhang et al., 2019). We recapitulated similar findings, and additionally observed an unexpected accumulation of TTR+ cells, suggesting a shift in fate or survival of neural precursors in MIC.

The cortical hem appears at embryonic day 8.5 in mouse, which will later differentiate into the non-neuronal epithelial cell component of the choroid plexus (ChP), expressing TTR, and serving as a midline telencephalic signaling center enriched for BMP and Wnt (Caronia-Brown et al., 2014; Dani et al., 2021; Parichha et al., 2022). Recent study implicates TTR+ cells in *in vivo* cortical growth rate, but why and how TTR+ cells expand at the expense of neurons, and whether patients with microcephaly patients have expansion of TTR+ cells remain open questions (Glass et al., 2025).

#### Polymicrogyria

Patients with polymicrogyria show excessive, irregular cortical folding. Because rodents lack cortical folding, modeling polymicrogyria in mouse has been difficult. Prior studies have uncovered several genetic causes, implicating primarily defective in neuronal migration (Akula *et al*., 2023; Epilepsy Phenome/Genome Project, 2021; Francis and Cappello, 2021; Jansen et al., 2016; Judkins et al., 2011; Smith *et al*., 2021; Stutterd and Leventer, 2014). Our results additionally implicate defective apical tight junctions, along with reduction of TBR2+ cells. Prior studies pointed to a critical role for tight junctions in maintenance of radial glial scaffolding (Akula *et al*., 2023; Giagtzoglou et al., 2009; Nakase and Naus, 2004; Wang *et al*., 2023a). Whether reduction of TBR2+ cells result from radial glial scaffolding defects could be investigate. We also found disparities between RNA and protein levels for TBR2, where transcript levels seemed comparable between controls and patients, but immunostaining showed substantially reduced and patchy expression (Figures 3A and G). These results may implicate alterations in protein translation or stability.

#### Epilepsy

Epilepsy, characterized by recurrent unprovoked seizures, is usually a result of altered excitation/inhibition ratio resulting in altered electrical discharges (ILAE, WHO, updated 2023). Numerous mechanisms have been proposed, including deficient GABAergic or excessive glutamatergic tone, altered channel properties, or astroglia synaptic regulation (Coulter and Steinhauser, 2015; Vezzani *et al*., 2022). Although synaptic physiology was not studied here, hBOs derived from patients with severe seizures showed excess astrogliosis, consistent with clinical observations (Rho and Boison, 2022). Analysis of neuroglial and synaptic functions could help uncover further mechanisms.

#### Intellectual disability

There are over 2000 single gene disorders that include intellectual disability as part of the clinical presentation, often associated with comorbid autism spectrum disorder, and implicating a diversity of genes. Cellular and transcriptome studies of hBOs in autism has documented biases in proportions of GABAergic and radial glia cells at DIV35 and DIV52 (Jourdon *et al*., 2023; Paulsen *et al*., 2022). Our study expands on these results and highlights involvement of non-neuronal cell types, for example TTR+ cells, suggesting altered neurotransmitter pathways. But how these cells contribute to the pathogenesis of intellectual disability remain elusive. The future might see these questions addressed with hBOs to further bridge cellular and neurocognitive defects.

## LIMITATIONS OF THE STUDY

Our study was limited to only four categories of neurodevelopmental disorders. Many other types of disease were not included. We studied predominantly families with recessive inheritance that transmitted biallelic pathogenic mutations, and while these demonstrated robust phenotypes at early timepoints in culture, confirmation of these results in other types of gene mutations is important. Our study relied on histopathology of organoids at early timepoints, so it is possible that other features might appear at later culture timepoints. Furthermore, a single IPSC clone was studied for each patient, so line-specific differences were not explored. Our CRISPR rescue experiments demonstrated a clear rescue for the microcephaly phenotype, but the less severe phenotype of intellectual disability limited analysis to gene expression. Future advances in stem cell models in combination with bioengineering or ‘assembloids’ might further improve on these hBO models.

**Table 1.**
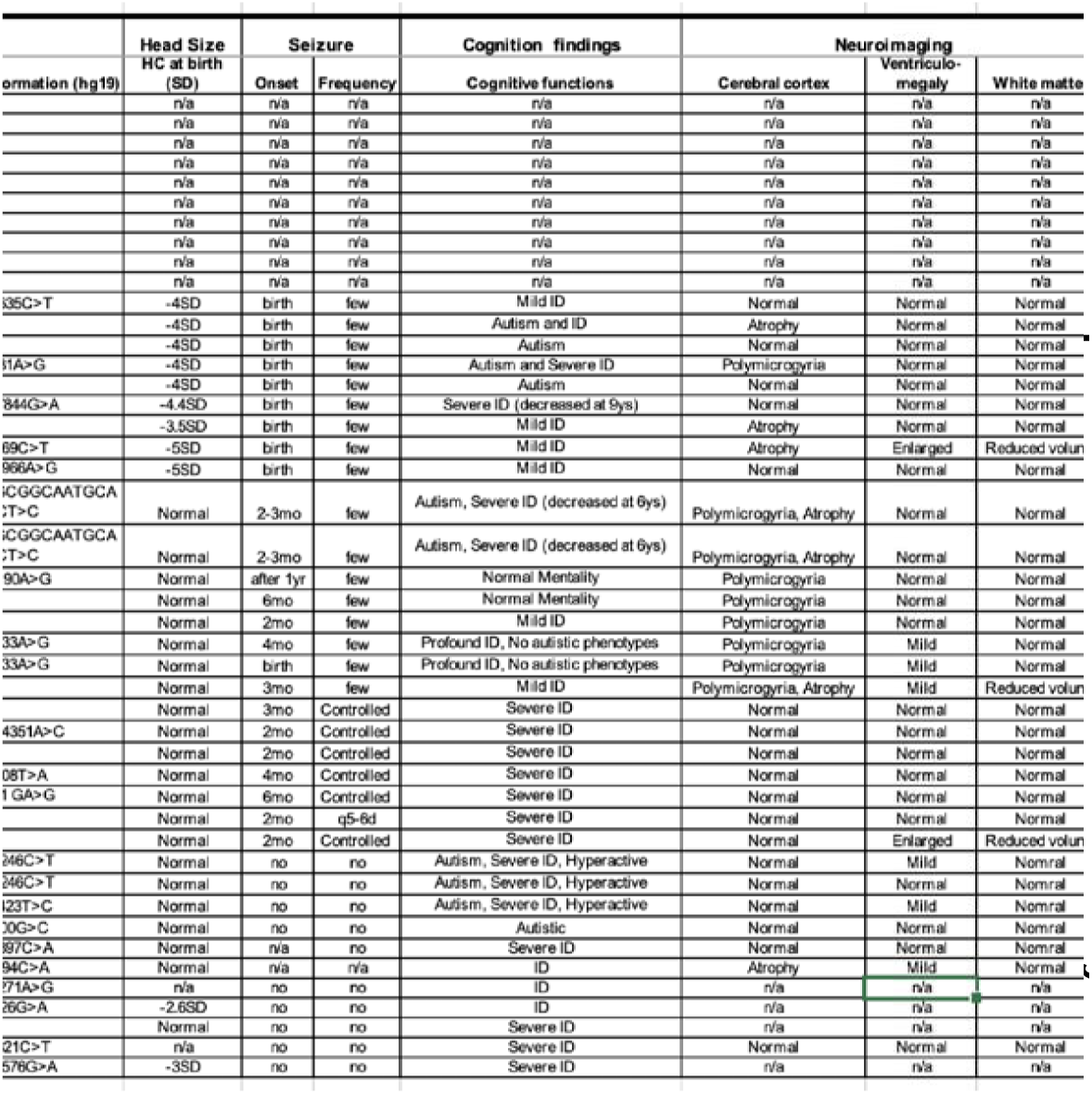
Clinical, genetic and hBOs information for patient’s iPSC lines in this study.

## ACKNOWLEDGEMENTS

We thank the subjects involved in this study. L.W. received support from NIH/NINDS Pathway to independent K99/R00 award (1K99NS125106-01A1, 4R00NS125106-03), CIRM Training Grant Postdoc award (EDUC4-12804); This work was supported by the Rady Children’s Hospital Neuroscience Endowment and HHMI to J.G.G. the UCSD Microscopy Core (NINDS grant P30NS047101), CIRM grant IT1-06611 for collection of patient cells and phenotypes, CIRM grant IR1-06600 to FujiFilm Cellular Dynamics, Inc for generation of IPCSs, the Yale, GMKF and NIH X01HG0113 for genotyping at CIDR, NIH X01HD100698 to support genome sequencing; The UCSD IGM Illumina NovaSeq 6000 was purchased with funding from NIH SIG (#S10 OD026929). We thank Drs. Lillia Lakoucheva, Uta Grieshammer and Larry Goldstein and anonymous reviewers for improvements on the manuscript.

## AUTHOR CONTRIBUTIONS

L.W. and J.G.G. conceived and designed the study. M.S.Z. recruited subjects. L.W. performed experiments with help from D.S., Y.N., Y.L., C.O.H., E.T., R.Y.Z., T.N., P.S., and K.C. Y.N. assembled clinical information with help from C.B., M.Y., and N.B.; J.L., J.W., R.H., and E.M. for data analysis. All authors reviewed and approved the manuscript.

## DECLARATION OF USE OF AI

Authors used ChatGPT to check grammar. The authors incorporated suggestions by hand when appropriate. Authors take full responsibility for the content of the publication.

## STAR METHOD

Detailed methods are provided in the online version of this paper and include the following:

- KEY RESOURCES TABLE
- **RESOURCE AVAILABILITY**

- Materials Availability
- Data and Code Availability
- EXPERIMENTAL MODEL AND SUBJECT DETAILS

- Human Subject
- Human Pluripotent Stem Cells
- METHOD DETAILS

- Human Pluripotent Stem Cells
- Human brain organoids (hBOs) Cultures
- Immunostaining and imaging for hBOs
- hBOs dissociation
- Single cell library preparation and sequencing
- scRNA-seq data preprocessing, clustering, and cell type annotation
- Differential cellular proportion test and pseudobulk differential expression test
- QUANTIFICATION AND STATISTICAL ANALYSIS

## RESOURCE AVAILABILITY

### Materials Availability

This study generated a library of genetically diverse iPSC lines from patients with NDDs. All iPSC lines are available in the Gleeson lab and can be shared upon reasonable request.

### Data Availability

The accession number for the sc-RNA-seq data reported in this study has been restored in PRJNA1088504 (Table S3).

https://dataview.ncbi.nlm.nih.gov/object/PRJNA1088504?reviewer=nrf2a0e8irmsnu6btmdofjco2e.

The whole genome or exome data for all iPSC lines are uploaded to dbGaP under phs002032, phs000288, phs003520, phs001267, phs001510, phs001822, phs001060, phs002621 accompanying phenotype data (https://www.ncbi.nlm.nih.gov/projects/gapprev/gap/cgi-bin/study.cgi?study_id=phs003520.v1.p1)(See Table S1 for details).

Patient brain images (MRI or CT) that correspond to each of the iPSC lines are available in Zip File 1 and https://brain-org-ndd.cells.ucsc.edu/.

All data and metadata are browsable via <https://brain-org-ndd.cells.ucsc.edu/>, facilitating reuse and future hypothesis generation.

Data supporting study findings are available from Lead Contact, Dr. Joseph Gleeson or upon request.

## EXPERIMENTAL MODEL AND SUBJECT DETAILS

### Human Subject

The study protocol was approved by the UC San Diego (UCSD) institutional review board protocol 140028. Informed consent was obtained from all participants or their legal guardians at the time of enrollment. Genome/exome data is available at dbGaP under documented accession numbers, including phs003520.v1.p1, see details in Table S1. All patient brain images are available at the UCSC cell browser at <https://brain-org-ndd.cells.ucsc.edu/>.

### Human Induced Pluripotent Stem Cells

Human induced pluripotent stem cell (iPSC) lines used in the current study were generated by CIRM (CIRM-IT1-06611); All hiPSC lines reprogrammed at Cellular Dynamics, Inc (Madison WI) and passed quality control measures for chromosomal integrity (by SNP microarray), pluripotency (by analysis of gene expression by qPCR of 48 mRNAs as well as immunostaining for OCT4 protein) and identity confirmation (by genotyping) and tested negative monthly for the presence of mycoplasma.

## METHOD DETAILS

### Human Pluripotent Stem Cells

H1 ESC (sex typed as male) were obtained from ATCC (CRL-11268^TM^) and WiCell (WAe001-A) and were maintained at low passage. Human induced pluripotent stem cells (hiPSCs) were from CIRM (CIRM-IT1-06611). All cells were regularly mycoplasma negative. H1 cells and hiPSCs were maintained in culture with mTeSR^TM^1 culture medium (STEMCELL Technologies) on Matrigel coated dishes (Corning).

### Human brain organoid (hBO) culture

H1 and hiPSCs were maintained in mTeSR and passaged according to manufacturer’s recommendations. Cortical brain organoids were generated as previously described (Kadoshima *et al*., 2013; Wang *et al*., 2020). H1 cells at 36 passages and hiPSC at 12 passages were dissociated into single cells with Accutase. In total, around 9000 cells were then plated in each well of an ultra-low-attachment 96-well U shaped plate (Corning, CLS3474) in “Cortical differentiation medium” with 20 µM Rock inhibitor (Selleckchem), 5 µM SB431542 (R&D), 3 µM endo-IWR-1 (Selleckchem) for the first 4d, then cultured for an additional 13d in cortical differentiation medium with 5 µM SB431542, 3 µM endo-IWR-1, with medium changes every 2d. On 18d, the immature organoids were transferred into ultra-low-attachment 6-well plate and cultured for another 16d in “Organoid differentiation medium.” Media was changed every 3–4d. On 35d, media was changed to “Maturation medium” supplemented with 1% Matrix High Concentration (HC), Growth Factor Reduced (GFR) Matrigel (Corning), with medium replaced every 3–4d for additional 34d. HC-GFR Matrigel was added fresh at each change. On 70d, media was changed to “Long-term maintenance medium” supplemented with 2% HC-GFR- Matrigel, 50 ng/ml BDNF (R&D, 248-BD-025/CF) and 50 ng/ml GDNF (R&D, 212-GD-010/CF) for long term organoid maintenance. “Cortical differentiation medium”: Glasgow’s MEM (GMEM), 20%Knock-Out Serum Replacement (KO-SR), 1X Non-Essential Amino Acid (NEAA), 1X Sodium Pyruvate, 1X β-Mercaptoethanol, 1X Pen/Strep. “Organoid differentiation medium”: DMEM/F12, 1X N2, 1X NEAA, 1X CD-lipid concentrator, 1XPen/Strep. “Maturation medium”: DMEM/F12, 10%FBS, 1X N2, 1X CD-lipid concentrator, 1X Pen/Strep, 1%HC-GFR Matrigel, 30μg/ml Heparin. “Long term maintenance medium”: DMEM/F12, 20%FBS, 1X N2, 1X B27 w/o vitamin A, 1X CD-lipid concentrator, 1X Pen/Strep, 2% HC-GFR Matrigel, 30μg/ml Heparin, 50ng/ml BDNF, 50ng/ml GDNF, 50ng/ml NT3.

### Immunostaining and imaging for hBOs and iPSCs

hBOs were fixed in 4% paraformaldehyde for 72h washed with PBST (PBS with 0.25% Tween 20) three times for 5 min, allowed to sink in 30% sucrose at least overnight at 4 °C, embedded in 15%/15% gelatin/sucrose solution and sectioned at 20 µm. 30min antigen retrieval (Tri- sodium citrate, 0.1% Tween-20, pH 6.0) was followed by 20min 0.5% Triton X-100 permeabilization at room temperature, then blocked with 5% BSA in PBS for 1 h at room temperature. After washing with PBST 3 times for 5 min, sections were incubated with primary antibodies in 5% BSA/Triton X-100 in PBS at the following dilutions: SOX2 (R&D, MAB2018R- SP, 1:100), TUJ1 (Biolegend, 801202, 1:1000), Cleaved Caspase 3 (Cell Signaling Technology, 9661S, 1:500), KI67 (BD-Biosciences, 550609, 1:1000), CTIP2 (Abcam, ab28448, 1:500), TBR2 (Abcam, EPR19012, 1:250), TBR1 (Abcam, ab31940, 1:250), GAD65 (Sigma-Aldrich, SAB4200232, 1:100), VGLUT1 (Thermo-Fisher Scientific, MA531373, 1:100), OCT4 (Biolegend, 653701, 1:400), GFAP (Abcam, ab4674, 1:250) overnight at 4 °C, washed three times with PBST for 10 min, then incubated with secondary antibodies (Alexa Fluor^TM^ 488 donkey anti- mouse lgG (HL+LL), AB_141607, 1:1000; Alexa Fluor^TM^ 594 donkey anti-rabbit lgG (HL+LL), AB_141637, 1:1000, Alexa Fluor^TM^ 594 donkey anti-rat lgG (HL+LL), AB_10561522, 1:1000, Alexa Fluor^TM^ 594 donkey anti-mouse lgG (HL+LL), AB_141633, 1:1000, Alexa Fluor^TM^ 488 donkey anti-rabbit lgG (HL+LL), AB_2535792, 1:1000, Alexa Fluor^TM^ 594 donkey anti-chicken lgY (HL+LL), A78951, 1:1000, Alexa Fluor^TM^ 488 goat anti-mouse lgG (HL+LL), A-11001, 1:1000, Alexa Fluor^TM^ 594 goat anti-rabbit lgG (HL+LL), A-11012, 1:1000) together with DAPI (ThermoFisher Scientific, D1306, 1:50000) for 2 h at room temperature, washed with PBST three times for 5 min, and mounted with Fluoromount-G® (Southern Biotech, 0100-01). Patient iPSCs were cultured on a Matrigel-coated glass cover-slip in mTeSR1^TM^ medium before harvested for immunostaining against OCT4, fixed with 4% PFA for 15 min at RT, permeabilized in 0.2% Triton X-100 for 15 min at RT, then blocked with 2.5% BAS + 2.5% goat serum in PBS for 1h at RT. These glass-coverslips were incubated with primary antibody of OCT4 in 2.5% BAS+2.5% goat serum + 0.01% Triton X-100 solution overnight at 4C, washed three times with PBST for 10 min/time, then incubated with secondary antibodies (Alexa Fluor^TM^ 488 donkey anti-mouse lgG (HL+LL), AB_141607, 1:1000) together with DAPI (ThermoFisher Scientific, D1306, 1:50000) for 2 h at room temperature, washed with PBST three times for 5 min, and mounted with Fluoromount-G® (Southern Biotech, 0100-01). Images were taken with ZEISS LSM880 Airyscan and Leica-SP8 (OCT4) with post-acquisition analysis in ImageJ-6. All stainings were performed using a streamlined workflow that fits for high-throughput screen (over 6000 hBOs were assayed), implementing an identical protocol, including batch control of media and reagents, antibodies and microscopy studies.

### CRISPR/Cas9 Genome Editing

Generation of rescue iPSC lines was carried out utilizing the CRISPR/Cas9 system, as described(Caillaud et al., 2022; Simkin *et al*., 2022). Briefly, equimolar chemically modified target-specific crRNA and trans-activating ATTO-550-labeled RNA were mixed, then Cas9 protein added. 0.61µM of RNP complex, 3.17 µg of GFP plasmid, and 3 µM of Single-stranded oligonucleotides (ssODNs) were delivered into iPSCs via electroporation using the Human Stem Cell Nucleofector Kit 2 (Lonza) and the Nucleofector II system (Amaxa Biosystems). A total of 2 × 10L cells were electroporated using the B-016 pulse program and cultured in mTeSR Plus medium supplemented with 10 µM Y27632 and 0.2 µM HDR enhancer (IDT) for 24 hours. Fluorescence-activated cell sorting (FACS) was performed 48 hours post-electroporation to isolate cells that were double positive for ATTO-550 and GFP. Approximately 5,000 sorted cells were plated onto 10-cm culture dishes for clonal isolation. Individual clones were picked up and expanded for subsequent genotyping and hBO generation at day 5. The ssODNs for *NARS1* correction introduced the NM_004539.4:c.50T>C correction along with the NM_004539.4:c.48C>T silent mutation. The ssODNs for *ACTL6B* correction introduced NM_016188.5:c.460T>C correction along with the NM_016188.5:c.465C>T silent mutation. Sanger sequencing confirmed both correction of one or both mutant alleles along with the silent mutation. Phenotyping of these lines was performed identically to other non-edited clones. All oligos were listed in Table S12.

### qPCR with hBOs

For RNA quantification, hBOs were washed twice with PBS and lysed in TRIzol. RNA was extracted using the QIAGEN RNA extraction kit. Then, 1 µg of RNA was used to generate complementary DNA with the SuperScript III First Strand Synthesis Kit (Invitrogen); 20Lng of cDNA was used to perform RT–qPCR with the iTaq Universal SYBR Green Supermix with different primers (Integrated DNA Technologies) at a final concentration of 100 nM for each primer using a Bio-Rad Laboratories Real-Time PCR System. All primers were listed in Table S12.

### hBOs dissociation

hBOs were dissociated using AccuMax (STEMCELL Technology) with 10µM Rock inhibitor, washed with PBS once, then changed to 500μl AccuMax. hBOs were incubated at 37°C for 20 min, with gentle trituration at 10min. 3ml Neuronal Medium was added into the dissociation system, and gentle trituration was performed to obtain a single cell suspension, centrifuged at 500g, RT for 10 min, supernatant removed, and cells re-suspended with 100µl of Neuronal Medium.

### Single cell library preparation and sequencing

10X sc-RNA-seq-3’-V3.1 kit (10X Genomics) was used to generate the GEM, cDNA and library were generated according to the manufacturer’s instructions (10X Genomics). Briefly, live cells were partitioned into nanoliter-scale Gel Bead-In-Emulsions (GEMs) with the 10x Chromium Controller (10X Genomics), 5000 cells were targeted. Upon cell lysis and dissolution of the Single Cell 3′-V3.1 Gel Bead within the droplet, primers containing an Illumina P7 and R2 sequence, a 14 bp 10XBarcode, a 10 bp randomer, and a poly-dT primer sequence were mixed with the cell lysates and bead-derived Master Mix. Barcoded, full-length cDNA from poly- adenylated mRNA was generated in each individual bead, then individual droplets were broken and homogenized before the remaining non-cDNA components were removed with silane magnetic beads (Invitrogen). Libraries were size-selected, and the R2, P5 and P7 sequences were added to each selected cDNA during end repair and adapter ligation. After Illumina bridge amplification of cDNA, each library was sequenced using the Novaseq6000 with PE150bp, around 20M∼100M reads were obtained for each sample.

### scRNA-seq data preprocessing, clustering, and cell type annotation

scRNA-seq reads were mapped to the human genome (GRCh38) with Cell Ranger (v7.1.0) (Zheng et al., 2017). The raw feature-by-droplet matrices were then applied to CellBender (v0.2.0) to distinguish positive cells and remove ambient RNA background (Fleming et al., 2023). Samples that retained no cells after CellBender (v0.2.0) were excluded from the downstream analysis. Doublet scores were computed with Scrublet (v0.2.3)(Wolock et al., 2019).

Cells passed the quality control metrics (genes/cell>500, mitochondrial reads<0.5%, doublet score<0.2) were applied for downstream analysis in Seurat (v4.3.0)(Hao et al., 2021). The top 3000 highly variable genes after normalization with Sctransform v2 were applied for principal component analysis (PCA), and the first 50 PCs were integrated with Harmony (v0.1.1) (Korsunsky et al., 2019). We removed the standard stressed cells by Gruffi(Vertesy et al., 2022). And total number of 155,323 cells were applied to Leiden clustering (resolution=2), and then were visualized with UMAP.

Cell type annotation was performed with the combination of marker genes identified "FindAllMarkers" with Seurat (Table S5) and labels transferred from a recently published cell atlas of the developing human brain at first-trimester (Braun *et al*., 2023; Nano *et al*., 2025). The cell-by-gene count table from the reference dataset was imported as a Seurat object normalized by Sctransform v2. The annotated labels from the reference (cluster IDs, brain regions, brain subdivisions, cell classes, and developmental ages) were transferred with “FindTransferAnchors” and “TransferData” (Table S5).

Cluster stability at the levels of cell subclass and cell type was evaluated using the “bootstrapStability” function the the R package “scran”. Bootstrapping was performed with 20 iterations, and the cluster stability was reported as the ratio of the adjusted observation pair counts for each pair of clusters by setting “mode=ratio”.

### Differential cellular proportion test and pseudobulk differential expression test

Propeller (implemented in the R package speckle, v0.99.7) was used to identify the differences in cellular proportions between disease-hBOs and Ctrl-hBOs with statistical analysis (Phipson et al., 2022). Proportions were logit transformed, significance was measured with the “propeller.ttest” function. Sex differences were added as a covariate to the linear model. Significant proportion differences were called with groups with FDR < 0.2 in either the logit transformed test, or the arcsine transformed test. Dreamlet (v0.99.6) applied to measure differential gene expression (DGE). In brief, the Seurat object was converted to a SingleCellExperiment object, then applied DGE tests on the pseudobulk count matrix generated from each cell subclass (Hoffman et al., 2023). Pseudobulk counts were aggregated by cell subclass and sample using the raw counts from the “RNA” slot in the original Seurat object. *DLX2*+ IPC, mixed neurons and mesoderm cells were removed from analysis due to insufficient number of cells across samples (at least 20 cells per sample in at least 10 samples across diagnosis groups, i.e. few if any *DLX2*+ cells in Ctrl group). Pseudobulk counts were normalized and applied to voom with Dream weights using the “processAssays” function with the following parameters: min.cells = 5, min.count = 5, min.samples = 4, min.prop = 0.3. DE tests were performed for each disease vs. control with a linear model with sex and batch added as covariates. The full lists of DE results are provided in Table S9 and Zip-file-DEs downsampled (Support File-Figures 3-5). Log_2_FC was calculated for all genes from Dreamlet test for each cell subclass, and each disease vs. control group, respectively, followed by Gene Set Enrichment Analysis (GSEA) with clusterProfiler (v4.4.4), with sets from Gene Ontology Biological Processes terms with more than 10 genes and less than 250 genes (Mootha et al., 2003; Subramanian et al., 2005; Wu et al., 2021; Yu et al., 2012).

Cell level differential expression analysis was performed with NEBULA (v1.5.4) in linear model mode (NEBULA-LN) (He et al., 2021), following similar filtering criteria used in pseudobulk analysis (clusters represented by ≥4 subjects and ≥5 cells per cluster, and genes expressed in ≥5 cells). NEBULA-LN models were fitted individually per cell subclass with fixed effects for disease, sex, and batch, with sample included as a random effect term by the models. Differential expression significance was determined using the BH procedure (FDR), with genes having FDR < 0.05 considered significant. To benchmark NEBULA-LN against the pseudobulk Dreamlet method, we computed the Pearson correlation and coefficient of determination (R²) between log2FC values obtained from each method for every disease-cell type combination. The empirical cumulative distribution functions of FDR values from NEBULA models using actual patient disease labels versus random shuffled disease labels were compared to assess the robustness of the differential expression signal (Table S10, Support File-Figures 6-8).

### Quantification and statistical analysis

Statistical analysis was performed in R v3.5.3. For each of the iPSC lines, n=12 organoids, including 3 biological replicates, and 4 technical replicates corresponding to each of the biological replicates. For quantification of immunostaining, each of the imaging fields was divided into 4 counting areas (25% of the whole image), the number of immune+ cells and DAPI+ cells were counted for each of counting area, average of the immune+ cells were calculated and normalized to average of DAPI+ cells to assess the percent of immune+ cells in image J. hBOs generated for each iPSC line were performed by two independent blinded researchers in parallel for reproductivity. In order to avoid p-value inflation, each of the counts for a single iPSC line were collapsed onto a single point (Figures 2-5, panels B-E). See quantification data for statistics in Tables S2, S6 and S7.

In the classifier, the proportions of cells expressing selected proteins as a percent of DAPI+ nuclei per imaging field in immunostained NDD-hBO sections at DIV28 (CC3, KI67, and SOX2) and DIV52 (TBR2, CC3, KI67, SOX2, and CTIP2) were capped at 1, averaged across replicates, and arcsine transformed. These features were used as input to train linear discriminant analysis (LDA) models separately for the two timepoints with leave-one-out cross-validation to classify the three diagnostic categories (C, NS, S), using the R package “caret”. The performances of the models were evaluated with receiver operating characteristic curves and AUC (area under the curve) using the “pROC” R package.

For the quantification of GFAP+ cells, all images were quantified using ImageJ software. GFAP channels were duplicated and passed through a contrast-limited adaptive histogram equalization (CLAHE) to better visualize positive staining. Images were manually thresholded and total GFAP+ area (including area fraction) was measured from the CLAHE-adjusted images. To assess the integrated density and astrocyte area, ROIs of GFAP+ astrocytes (with a clearly defined nuclear signal) from the CLAHE-adjusted thresholds were defined and subsequently overlaid on the raw images. Statistical analysis was performed using GraphPad (Prism10). Data was tested for normality prior to analysis to ensure correct statistical tests were used.

## KEY RESOURCES TABLE

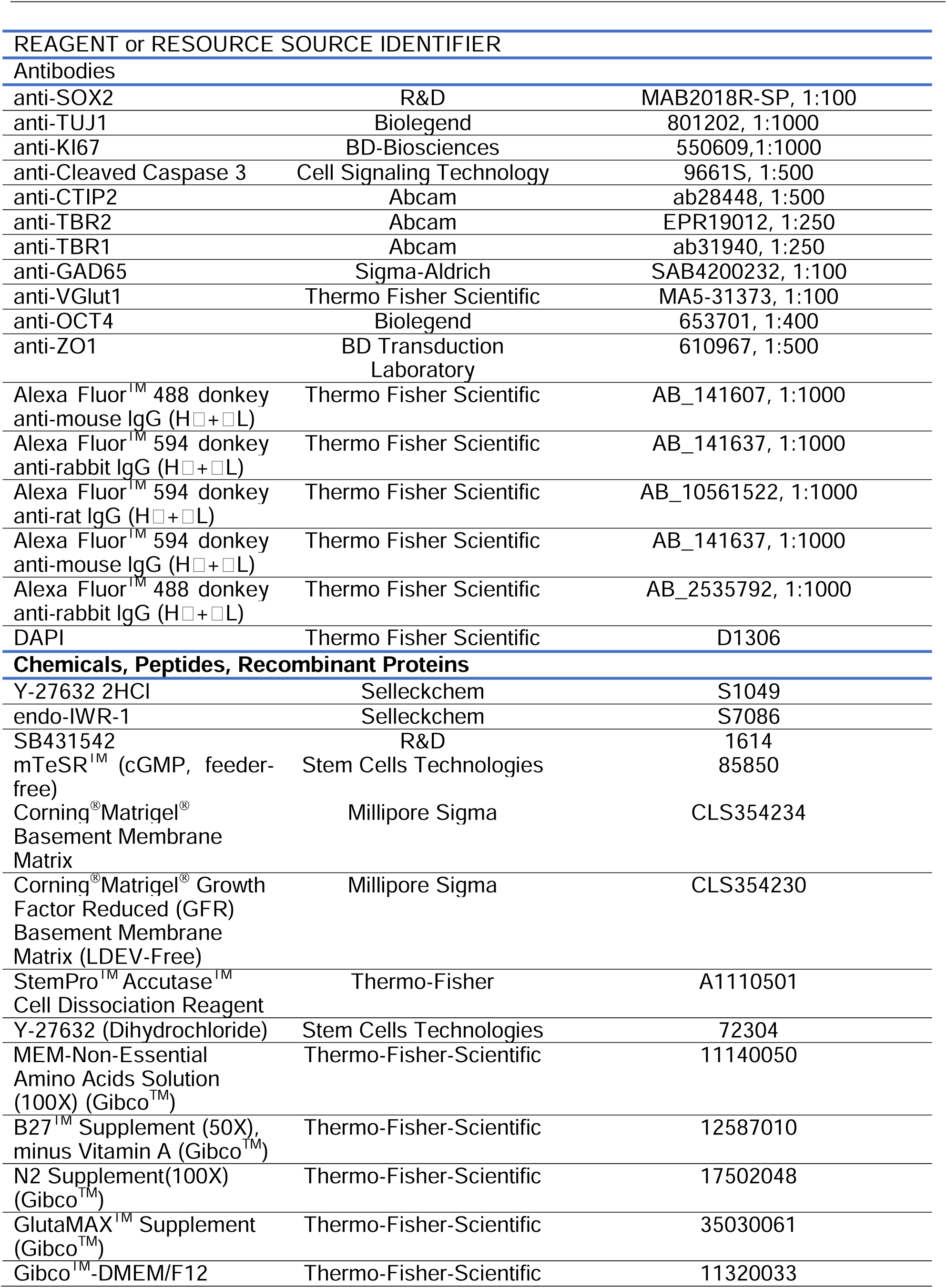
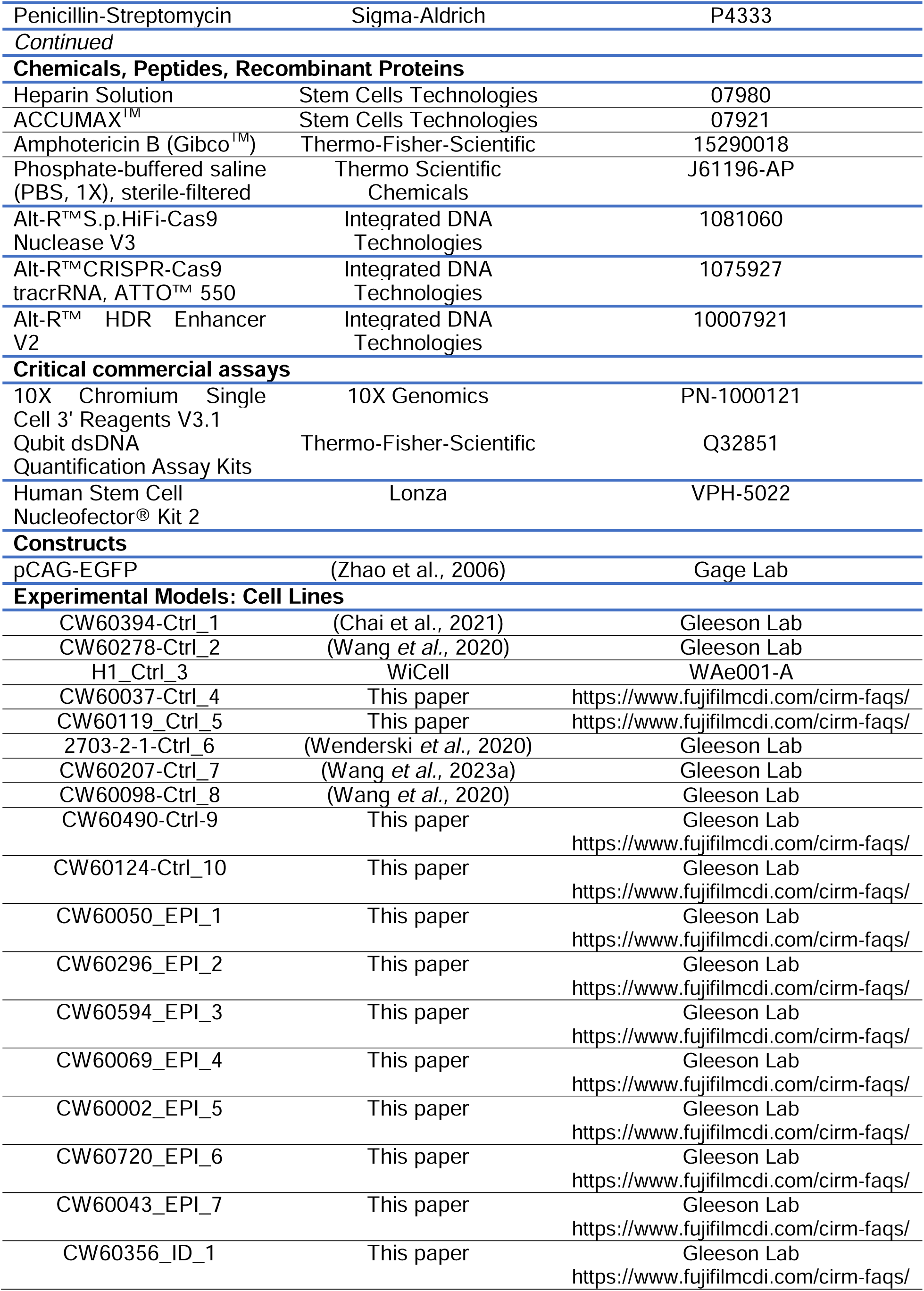
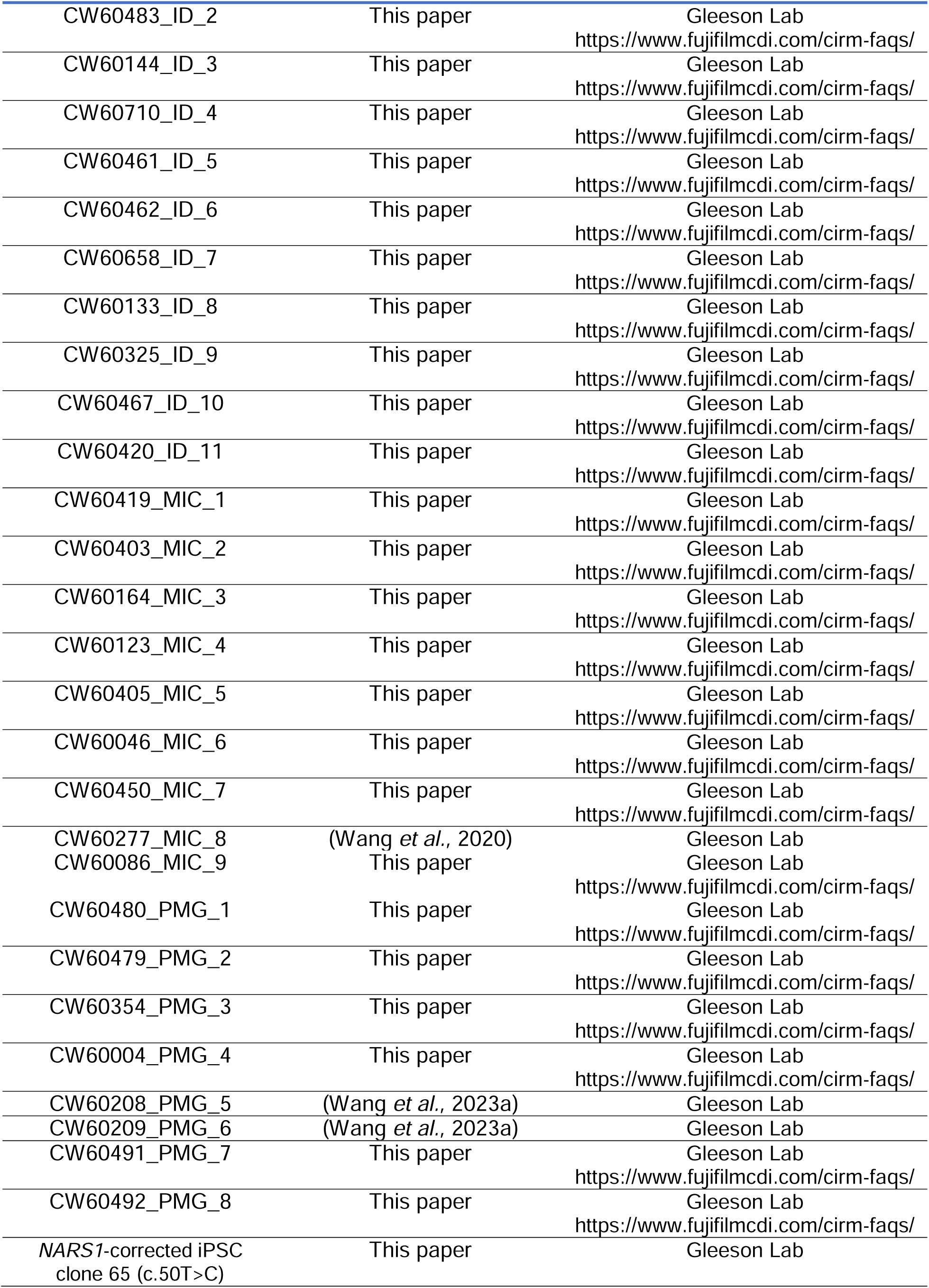
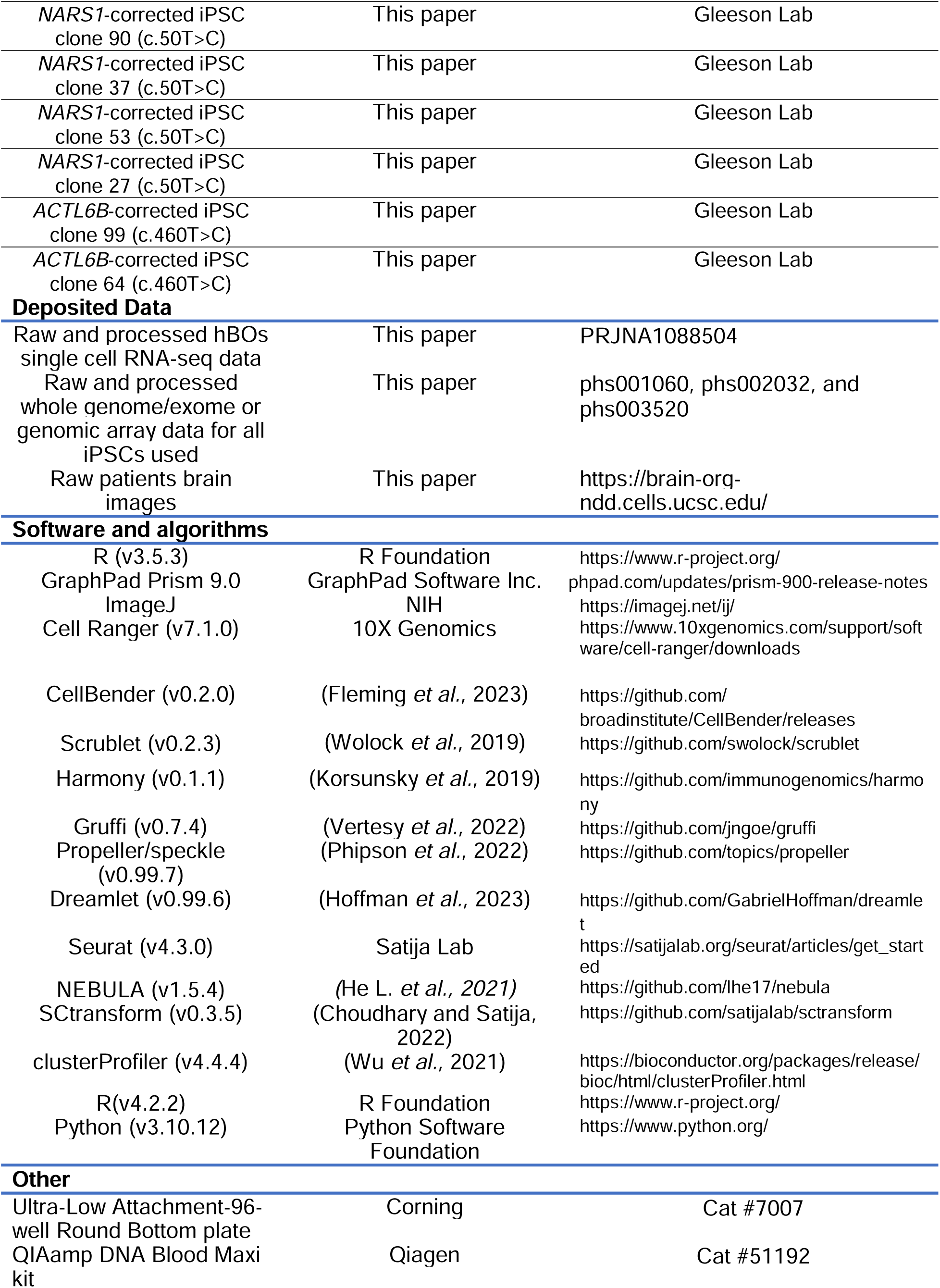

## Supplementary Figures

**Figure S1 related to Figures 1-5.**
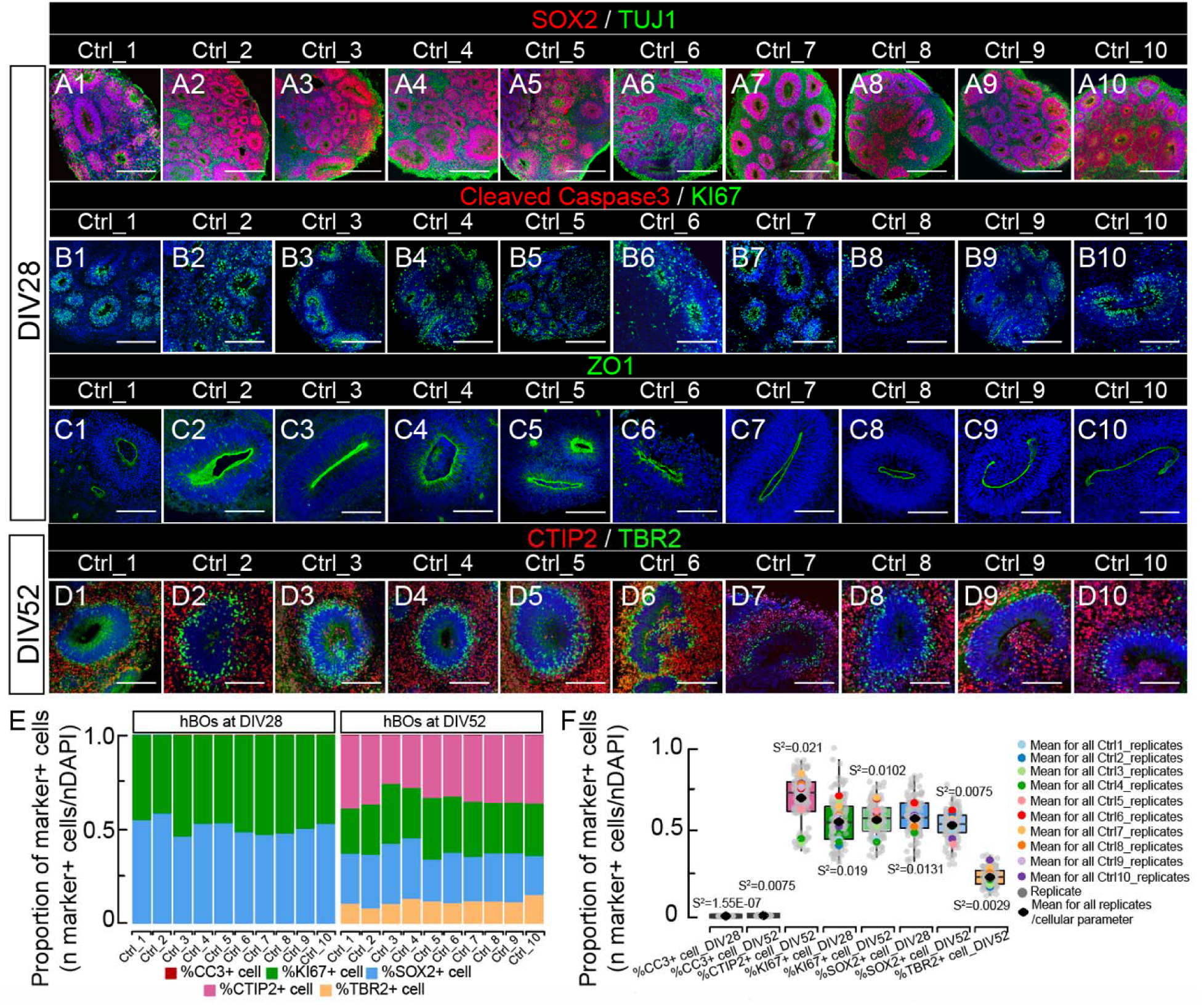
hBOs exhibited reproducible cellular features across iPSC lines. (A-D) 2D-immunostaining across ten healthy individual iPSC lines (Ctrl groups) showed similar histological features. Ctrl-hBOs were harvested for immunostaining for SOX2+ radial glial cells and TUJ1+ immature neurons (A1-A10), CC3 for apoptosis and KI67 for proliferation (B1-B10), ZO1 for apical tight junction (C1-C10) at DIV28, CTIP2 for deeper layer neurons and TBR2 for intermediate progenitor cells at DIV52 (D1-D10). DAPI: nucleus. Bar: 400mm. (E) Variability of cellular parameters across ten individual iPSC lines, based on numbers of positive cells per cellular marker. Percentage of marker-positive cells calculated for hBOs at DIV 28 and 52 for each replicate (technical and biological). Mean calculated per individual iPSC line, plotted using the mean. Raw quantification provided in Table S2-sheet 1. (F) Variation per cellular parameter across iPSC lines. Small gray dots: value for each replicate. Big solid dots: mean for all replicates per iPSC line. Rhombus: mean for all replicates per cellular parameter. Sample variance presented as S^2^.

**Figure S2 related to Figure 1.**
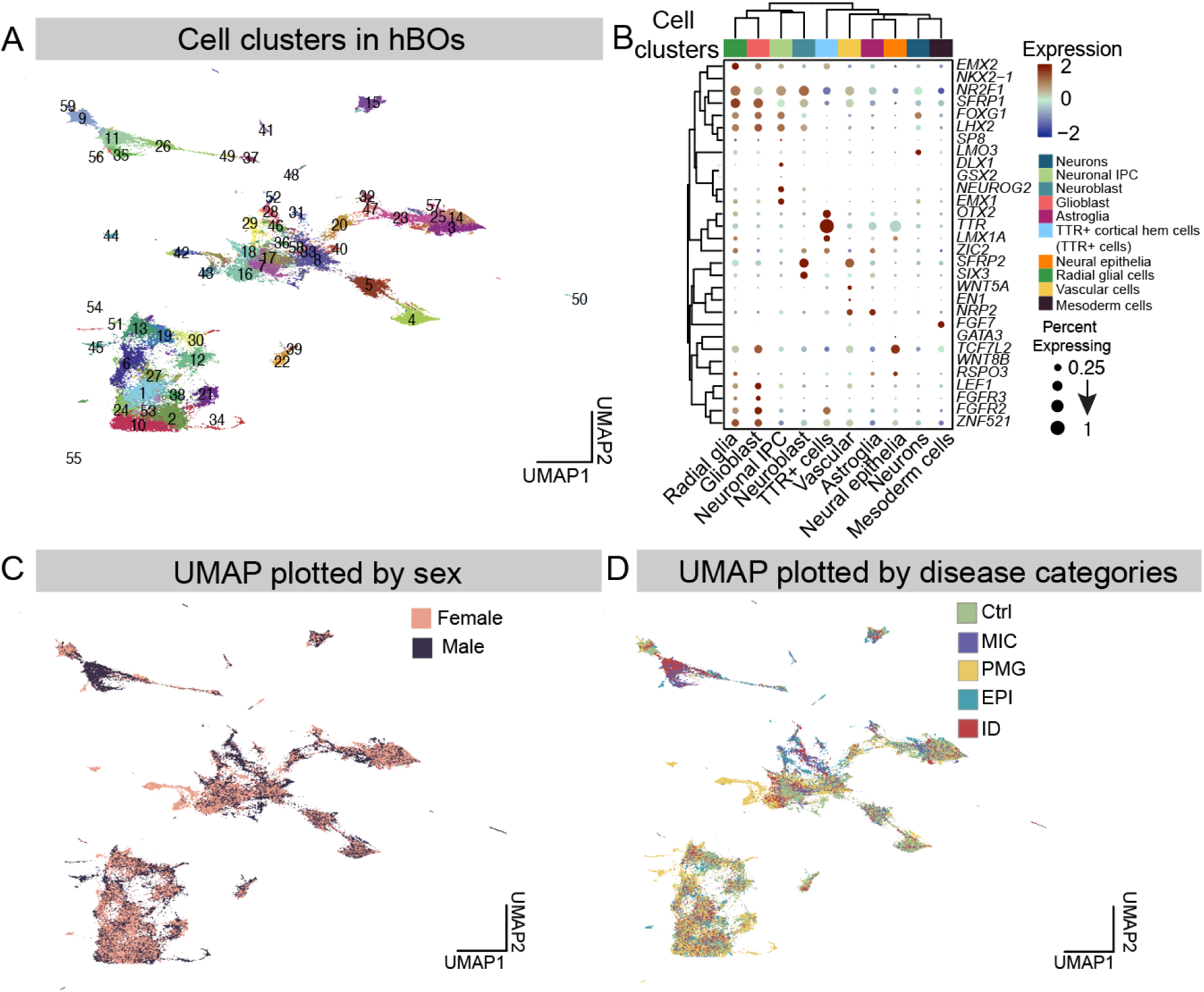
Sample quality from scRNA-seq analysis. (A) UMAP of all cellular clusters identified with scRNA-seq. (B) Specific gene expression for each of the major cellular classes. For instance, *EMX2* was highly expressed in radial glial cells (first column, top left) as expected. (C) UMAP plotted by sex of the sample analyzed. Note few male-female differences. (D) UMAP plotted by disease categories.

**Figure S3 related to Figure 1.**
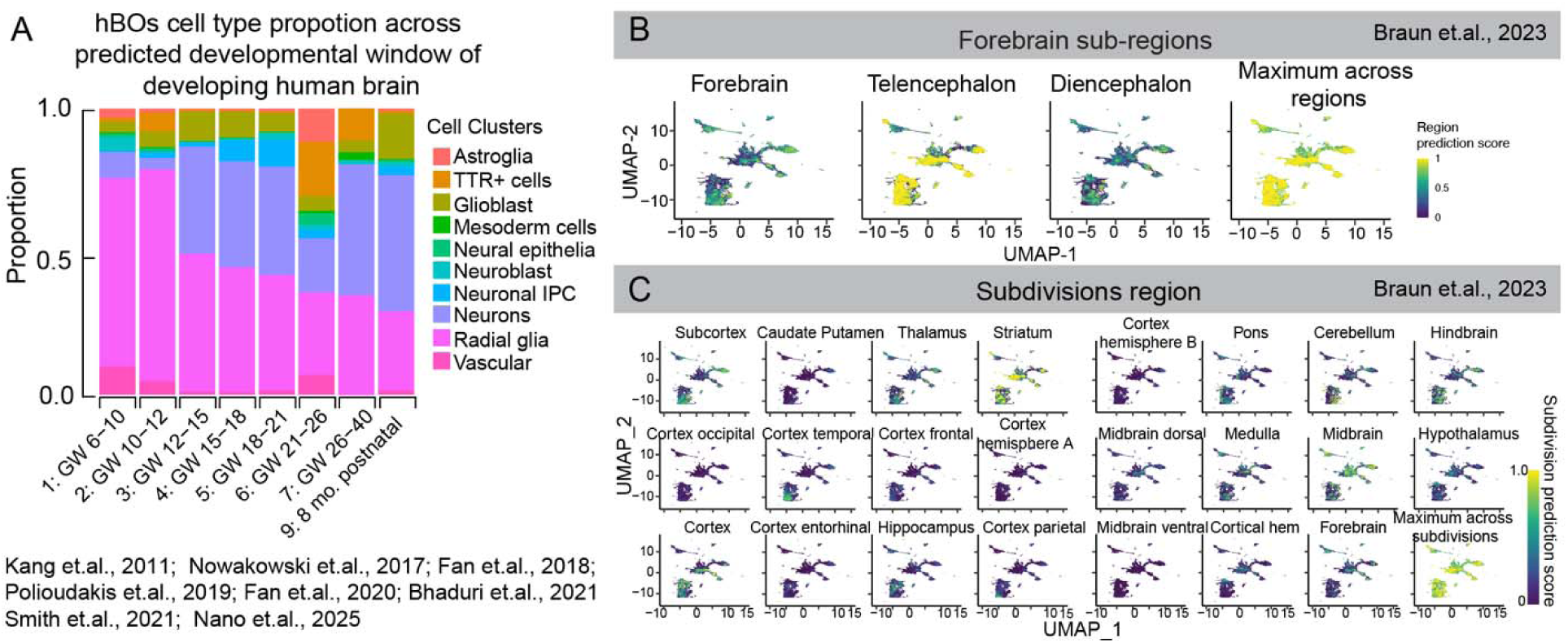
hBOs captured temporal and spatial features of developing human brain. (A) Distributions of the major cell types from hBOs mapped onto human fetal single cell data from publications listed. (B) Prediction for forebrain sub-regions in the developing human brain. Note telencephalon yields the highest prediction score. (C) Prediction for sub-regions in the developing human brain. Note striatum yielded the highest prediction score.

**Figure S4 related to Figure 1.**
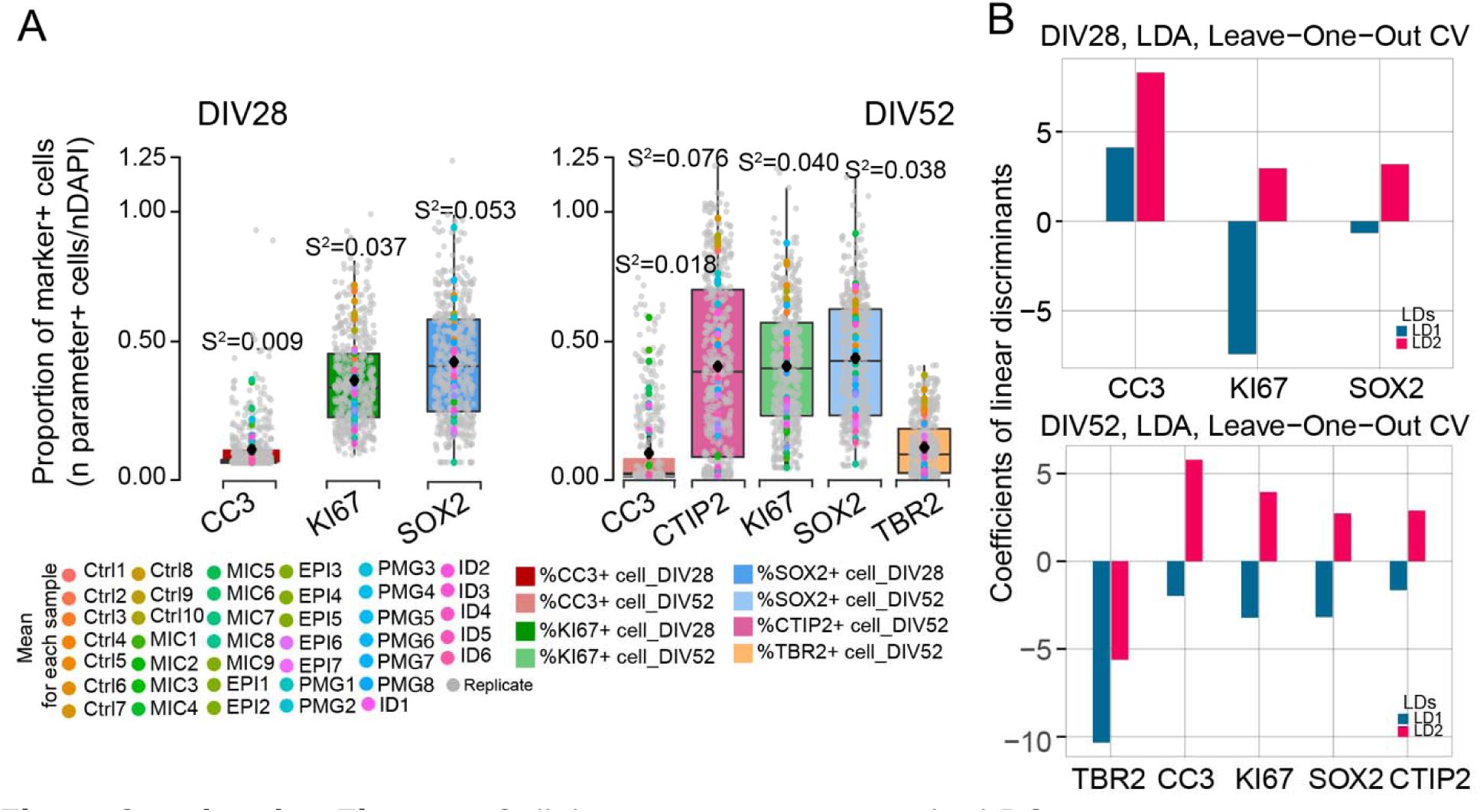
Cellular parameters across the hBOs. (A) Variation across all hBOs assayed for each cellular marker. Small gray dot: each replicate. Solid colored dot: mean for each sample. Rhombus: mean for all replicates per cellular parameter. Sample variances presented with S^2^. (B) Importance of markers as classifiers, represented as coefficients of the linear discriminants (LDs) for DIV28 and DIV52. LDA: linear discriminant analysis. CV: cross-validation.

**Figure S5 related to Figure 2.**
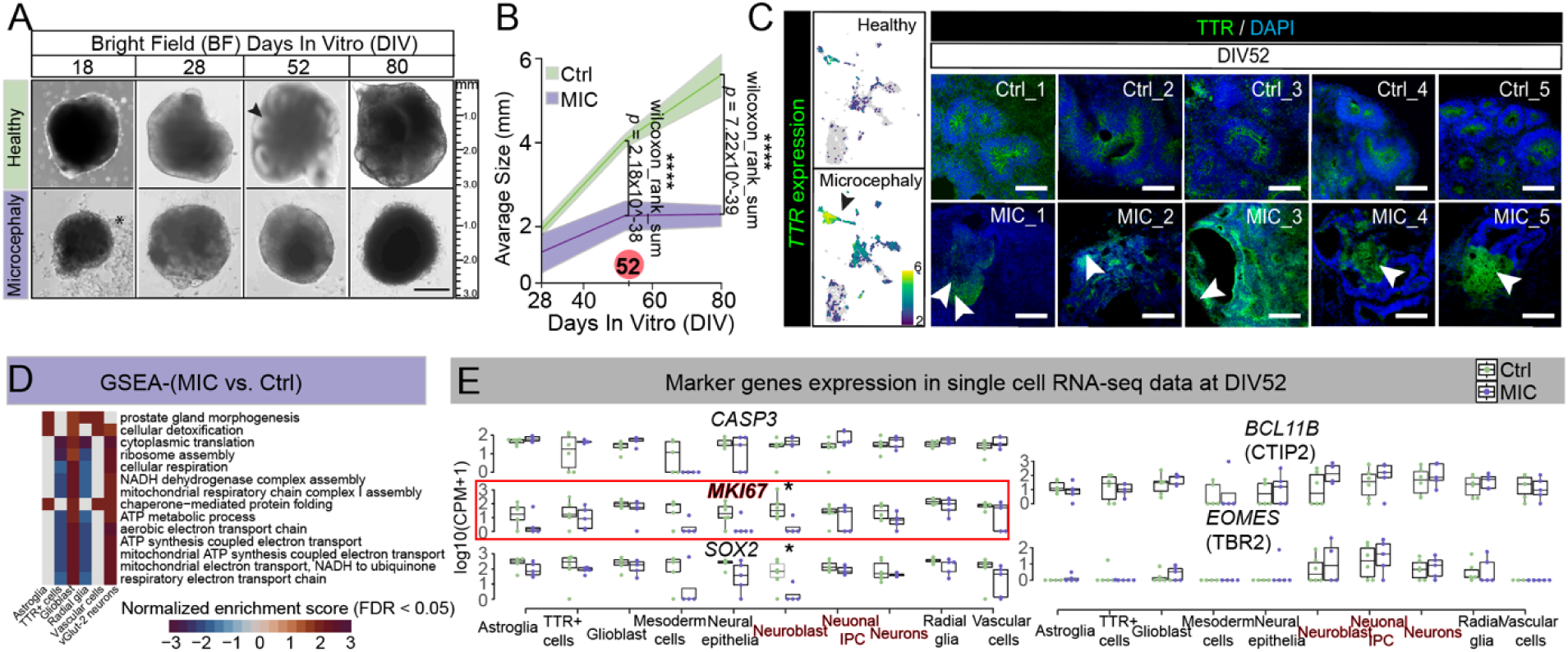
smaller hBOs and alterations in the transcriptome in microcephaly. (A) Bright field image showed smaller hBOs in MIC vs. Ctrl across timepoints. Note size of ctrl ranged from 2mm to 5mm in DIV28 to DIV52. Bar:1mm. Arrow: lobe with lumen like structure. Star: cell debris. (B) Quantification and statistics for the sizes of hBOs. ****p<0.00001. (C) Enriched of TTR+ cells (green) in MIC compared to Ctrl. DAPI: Nucleus; Bar: 400μm (D) GSEA with all genes expressed from pseudobulk analysis on MIC vs. Ctrl, normalized enrichment score calculated to reflect the enrichment, FDR<0.05. (E) Maker genes expression in different cellular clusters of the single cell RNA-seq results, * p<0.05 (Table S5).

**Figure S6 related to Figure 3-5.**
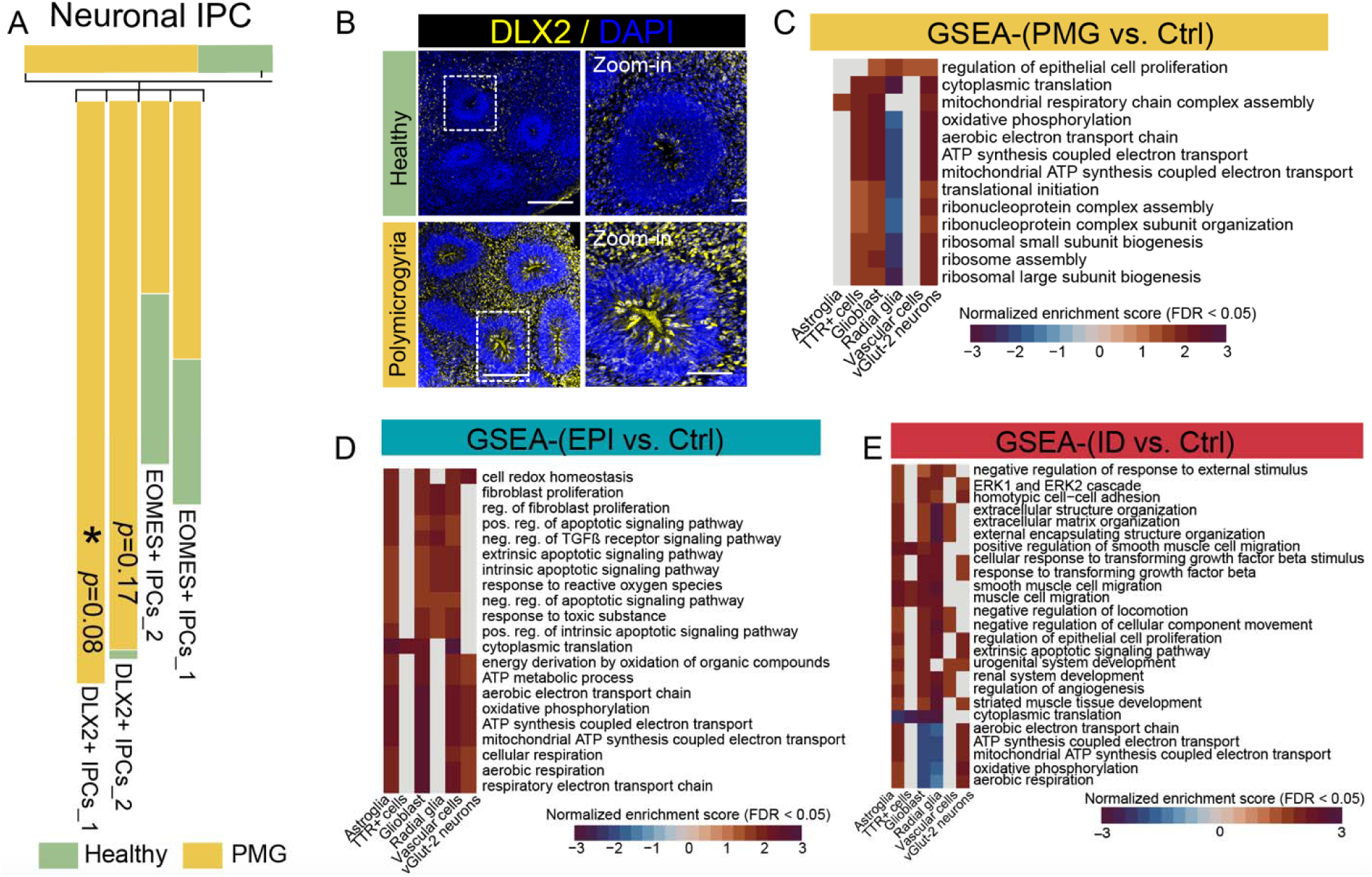
scRNA-seq for PMG, EPI, and ID. (A) Sub-clustering of neuronal intermedite progentior cells (IPSc) showed significant increase of the DLX2+ cells, but no significant changes observed for the EOMES+ cells in PMG. * p<0.1. (B) DLX2 (yellow) cells were increased in both signal intensity and number of cells in PMG at DIV52 section. DAPI: nucleus (blue), bar: 400μm, zoom-in from the white dash box, bar: 200μm. (C) Gene set enrichment analysis (GSEA) with all genes expressed from pseudobulk analysis on PMG vs. Ctrl, normalized enrichment score reflected enrichment, FDR<0.05. (D) GSEA with all genes expressed from pseudobulk analysis on EPI vs. Ctrl, normalized enrichment score reflected enrichment, FDR<0.05. (E) Differentially expressed genes in EPI vs. Ctrl in astroglia, glioblast, radial glia and vascular cells, Z-scaled expression by row based on level of expression, FDR<0.1. (F) GSEA with all genes expressed from pseudobulk analysis on ID vs. Ctrl, normalized enrichment score reflects enrichment, FDR<0.05.

**Figure S7 related to Figure 4.**
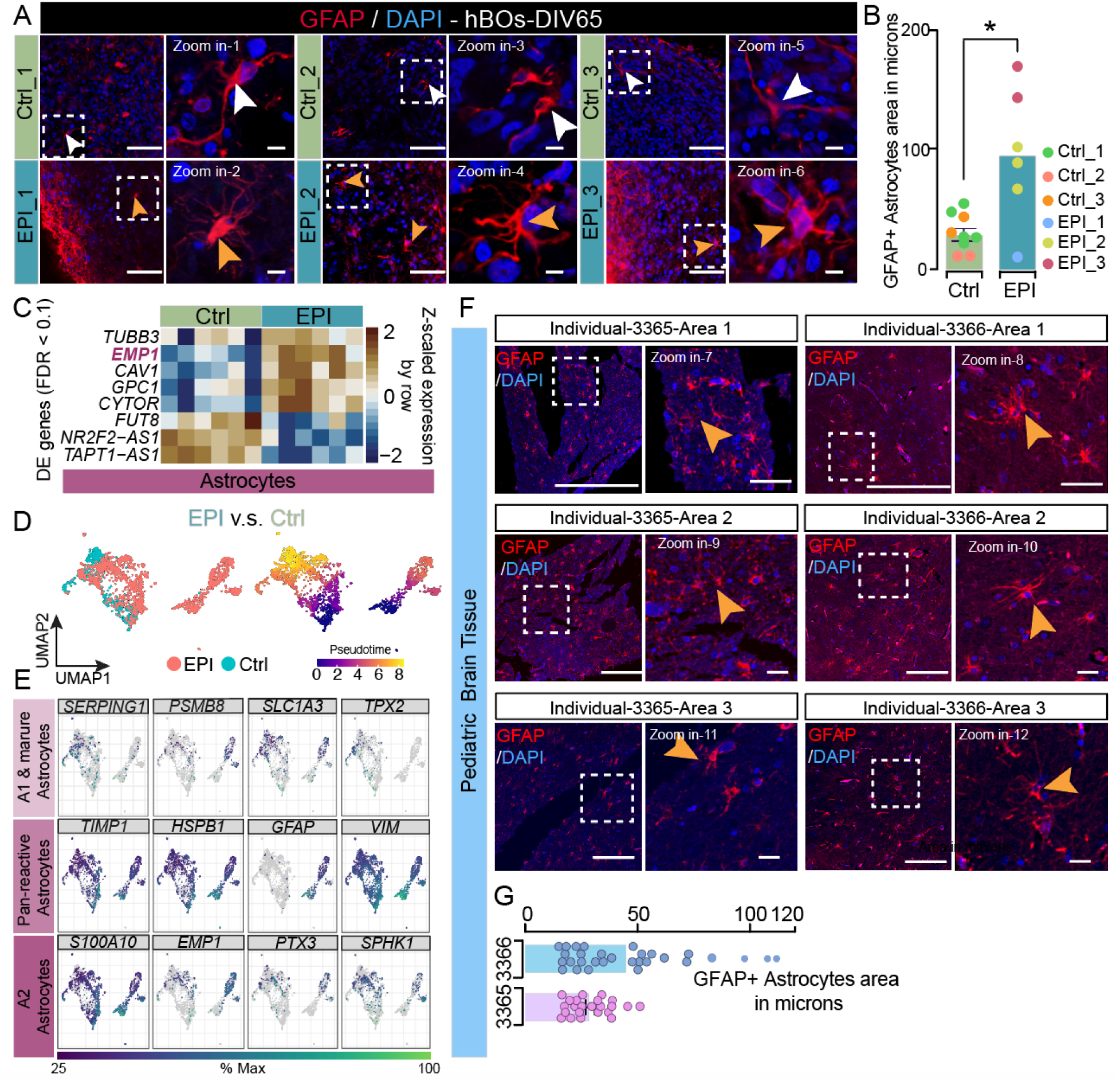
astrogliosis in epilepsy brain sections and hBOs from patients with epilepsy. (A) 2D-immunostaining for GFAP across Ctrl and EPI showed slight increase in astrocyte number as well as activated astrocytic morphology (Zoom-in view), suggesting astrogliosis. hBOs derived from three individual with epilepsy were assayed, respectively. White arrows: astrocytes in control; Orange arrows: astrocytes in EPI. DAPI: nucleus. Bar: 400μm. Zoom-in bar: 10μm. (B) Areas of GFAP+ signal (in microns), GFAP+ astrocytes manually counted. Significance with two-tailed p-value. \**p*<0.05, Mann-Whitney test. (C) Differentially expressed (DE) genes analysis in EPI vs. Ctrl in astroglia highlights *EMP1* increased expression. Z-scaled expression by row, FDR<0.1. (D) Cellular distribution in EPI and Ctrl. Left: merged UMAP for Ctrl and EPI. Right: pseudotime of predicted cellular trajectories. (E) Expression of specific astrocytic genes: Astrocytes termed ‘A1’ and ‘mature’ *SERPING1, PSMB8, SLC1A3*, and *TPX2*; Pan-reactive astrocytes genes: *TIMP1, HSPB1, GFAP,* and *VIM*; A2-astrocytes genes: *S100A10, EMP1, PTX3*, and *SPHK1* as previously reported *(Liddelow et al., 2017)*. (F) GFAP immunostaining in brain sections from epilepsy patients showed astrocytic activation represented by the morphologies of astrocytes. Three image areas presented for individual 3366 and 3365 (See details in Support File_Clinical Notes). DAPI: nucleus; Bar: 400μm. Zoom-in bar: 20μm. orange arrows: astrocytes. (G) Quantification of the images shown in F. Areas of GFAP+ signal was selected in microns, GFAP+ astrocytes were then manually selected and counted.

**Figure S8 related to Figure 5.**
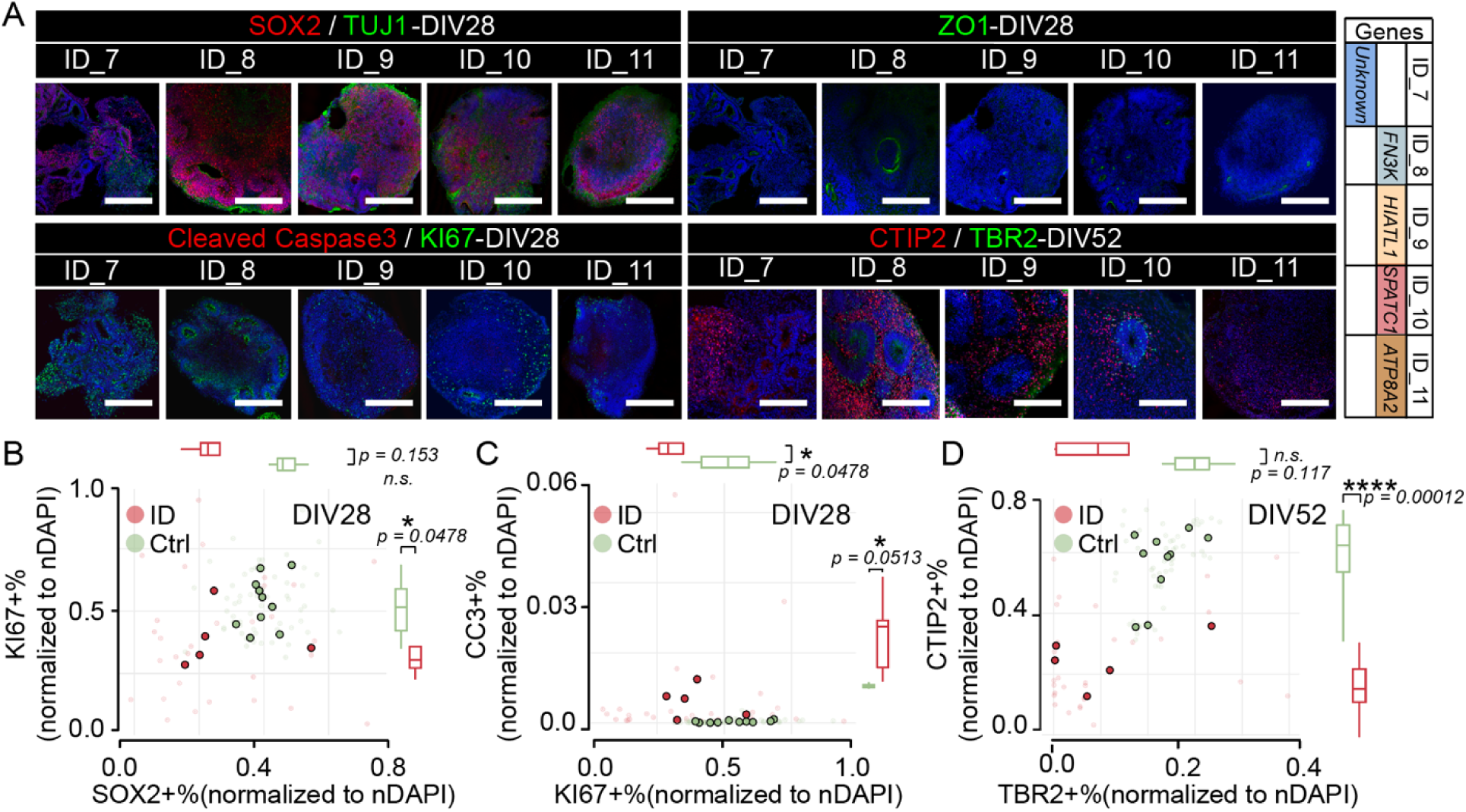
additional hBOs from patients with intellectual disability showed consistent cellular phenotypes. (A) Cellular immunostaining phenotypes in ID. hBOs derived from five patients at DIV28 and DIV52, four with established genetic cause (noted at right, e.g., *FN3K*). Comparison with 10 Ctrl (Fig. S1) shows apparent differences in neural rosette composition at DIV28 and scattered CTIP2+ and TBR2+ at DIV52. Bar: 400μm. (B-D) Scatter plots: significantly reduced KI67+ cells at DIV28 (B and C). No significant alterations were observed for SOX2 at DIV28 (B). C: increased CC3+ cells in ID-hBOs at DIV28. D: significant reduction in CTIP2+ and TBR2+ cells at DIV52. Two-sided t-test, *p<0.1, **p<0.01, ***p<0.001, and ****p<0.0001. Smaller dots: percentage of marker positive cells for each organoid derived from the iPSC lines assayed (ten individual control lines and five individual ID lines). Larger dots: mean of the percentage-positive cells from all organoids derived from the same iPSC line. *p* values: differences among individual iPSC lines (See detailed in Table S2-Figure S7B-D).

**Figure S9 related to Figure 2.**
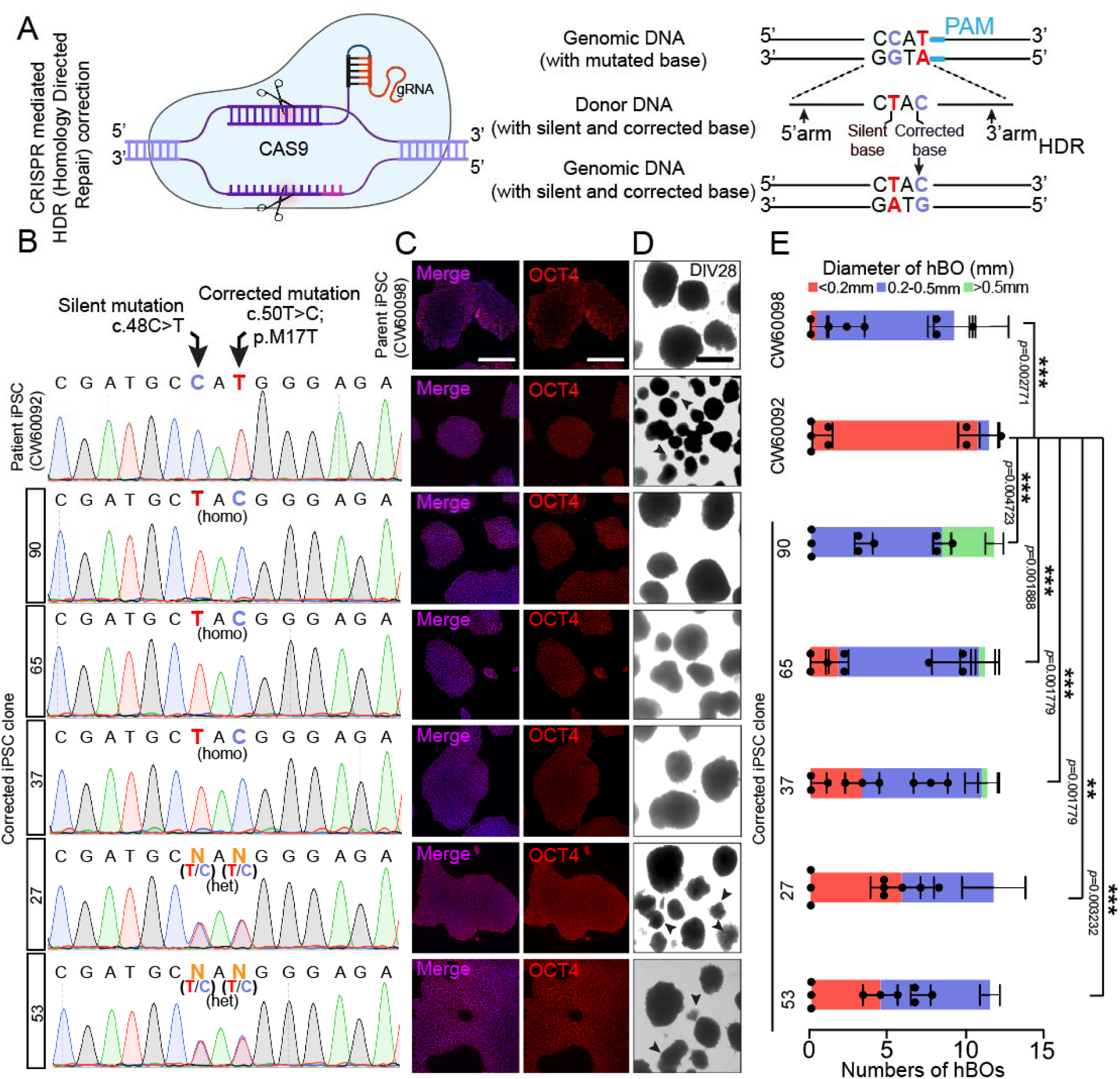
CRISPR mediated correction of *NARS1* mutation rescued patient hBO phenotype. (A) CRISPR mediated HDR correction of the patient mutation in *NARS1* gene. A silent mutation was introduced adjacent to the correction, to exclude the possibility that a deletion occurred at the locus. (B) Sanger sequencing: mutated (c.50T) and corrected (c.50T>C) base across multiple iPSC clones. A one-base silent mutation (c.48) was introduced. (C) OCT4 staining across the parent (CW60098), patient (CW60092), and corrected clones (90, 65, 37 for homozygous; and 53 and 27 for heterozygous) confirm pluripotency. DAPI: nucleus (blue). Bar: 400μm. (D) Bright images of hBOs derived from parent, patient, and corrected patient iPSC clones. hBOs derived from the corrected clones show comparable hBO size to the parent. Bar: 400μm. (E) Quantification for D. The size of hBOs from three independent biological replicates. *p<0.1, **p<0.01, ***p<0.001, and ****p<0.0001 (unpaired t-test with Welch correction).

**Figure S10 related to Figure 4 and 5.**
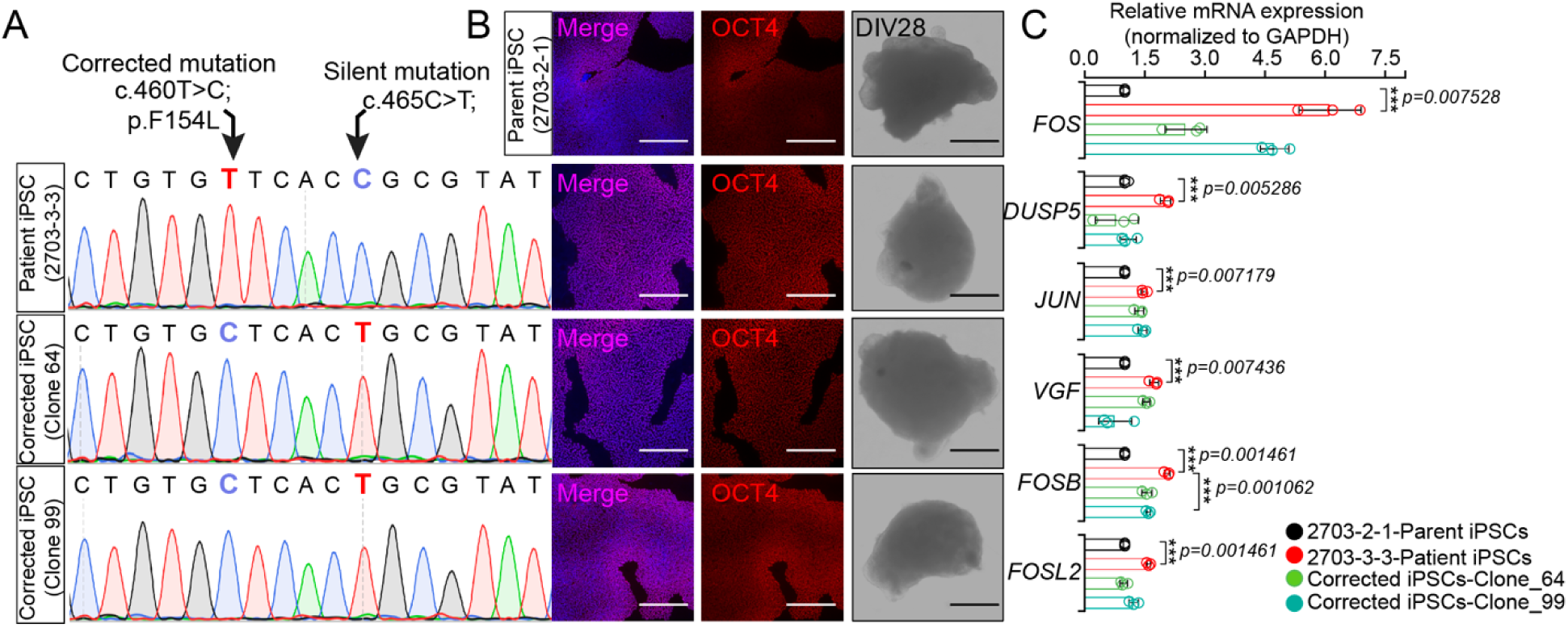
CRISPR mediated *ACTL6B* point mutation correction rescues up-regulation early-response genes in patient hBOs. (A) Sanger sequencing of mutated and corrected (c.460T>C) base across multiple iPSC clones. A silent mutation was introduced adjacent to the correction, to exclude the possibility that a deletion occurred at the locus. (C) Pluripotency for the parent (2703-2-1), patient (2703-3-3), and corrected clones (64 and 99) iPSCs lines evidenced by OCT4 staining. DAPI: nucleus (blue). Bar: 400μm. (D) No significant gross phenotypes were observed among hBOs derived from the parent, patient, and corrected iPSC clones. Bright images of hBOs derived from parent, patient, and corrected patient iPSC clones. (E) Levels of mRNA expression of the early response genes were partially rescued in hBOs derived from corrected clones. hBOs harvested at DIV28 to detect the mRNA expression of the early response genes, *FOS, DUSP5, JUN, VGF, FOSB*, and *FOSL2*. Three hBOs were applied as three independent biological replicates. Multiple unpaired t-tests were applied to measure the significance. Two-stage step-up method of Benjamini, Krieger and Yekutieli was used for multiple comparisons correction, FDR < 0.01. *p<0.1, **p<0.01, ***p<0.001, and ****p<0.0001.

## Support File Figures

**Support File-Figure 1, related to Figure 1-5.**
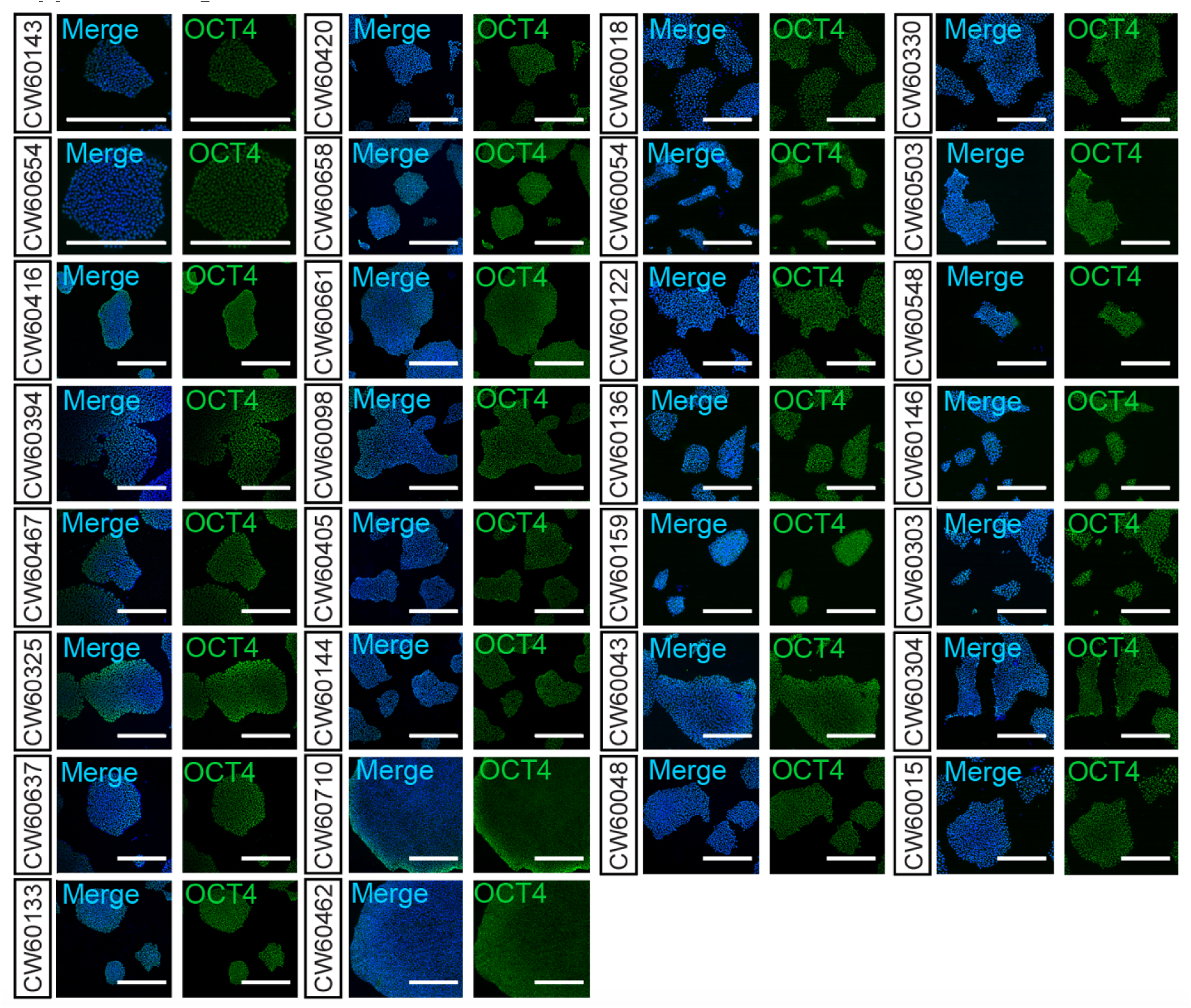
Left: OCT4 staining of patient iPSCs used in the study confirmed pluripotency. Right: additional lines from 14 randomly selected patients of the total 352 lines similarly cultured also showed positive OCT4 staining. DAPI: nucleus in blue. Bar: 400μm.

**Support File-Figure 2, related to Figures 1-5 and S1.**
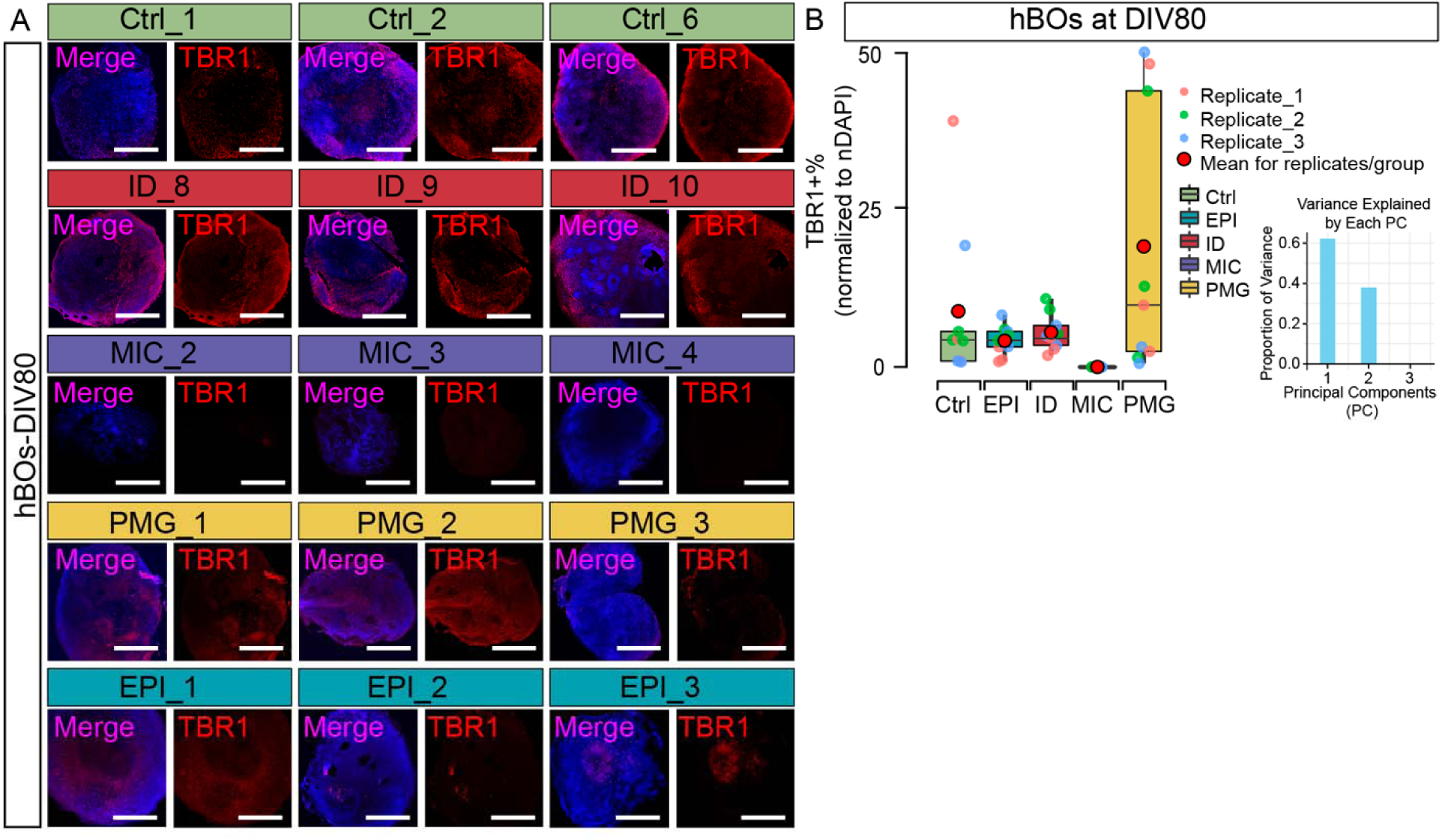
phenotype of hBOs at DIV80. (A) Immunostaining for TBR1 across NDD disease categories. DAPI: nucleus in blue. Bar: 400μm. (B) Percentage of TBR1+ cells across NDD assayed in A. Three hBOs from three independent biological replicates.

**Support File-Figure_3 related to Figure 1-5.**
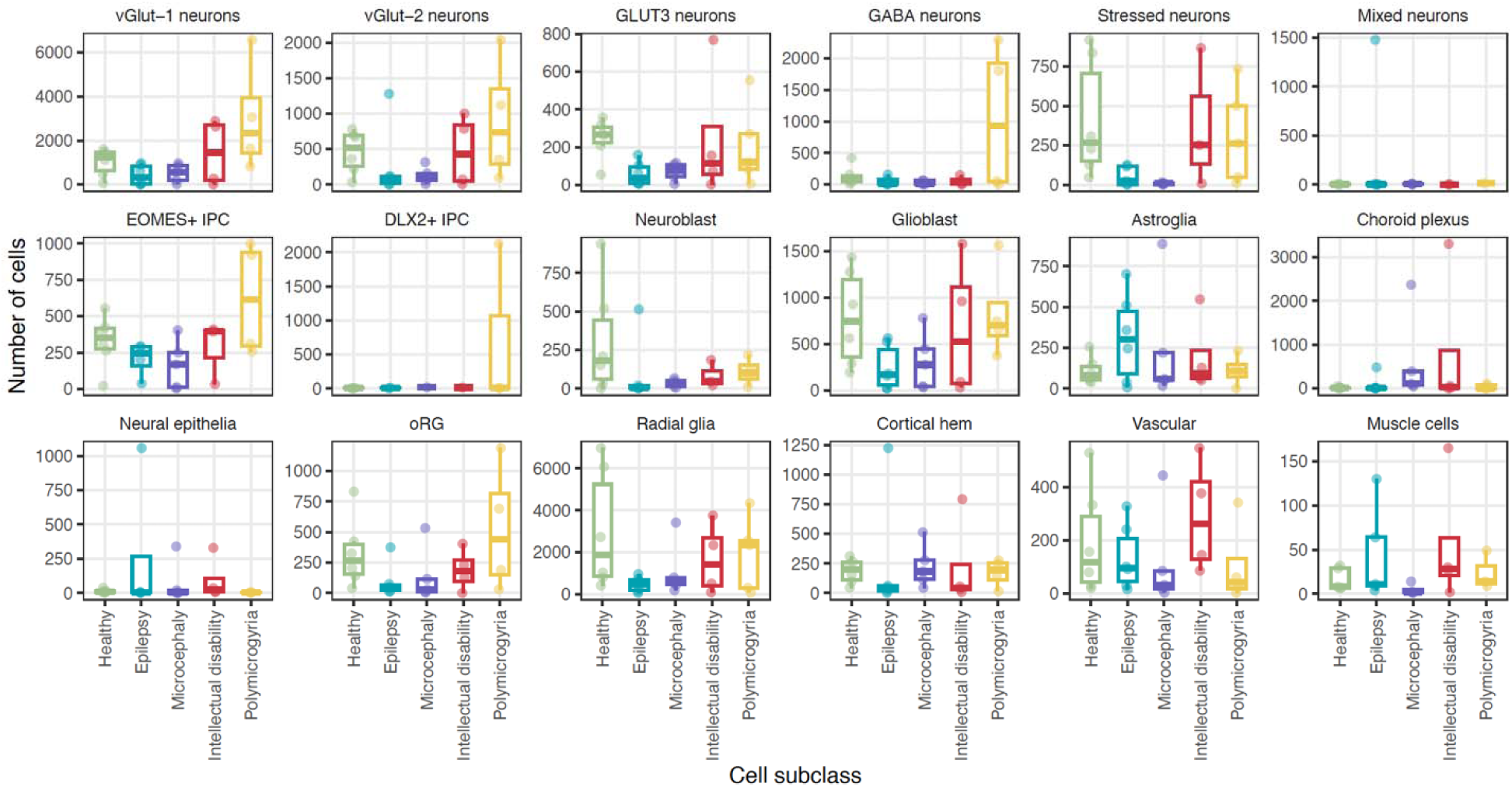
number of cells in each of the cell types of NDD- hBOs. Box plots distribution of the number of cells in each sample, grouped by cell subclasses and diagnosis. Lower and upper hinges correspond to the first and third quartiles; whiskers extend 1.5 * IQR (interquartile range) away from the hinges; center denotes median.

**Support File-Figure_4 related to Figure 1-5.**
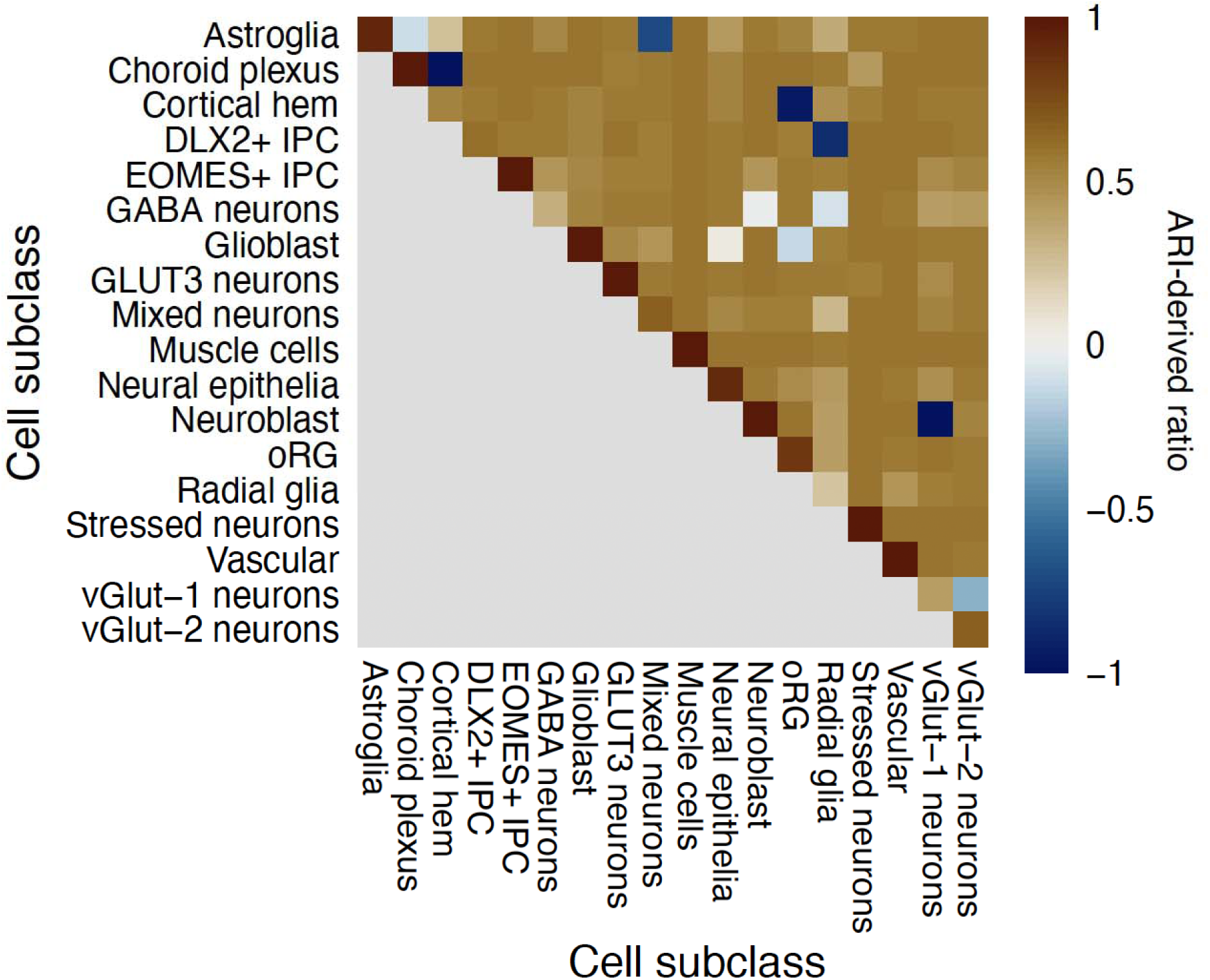
Cell subclass evaluated using bootstrapping with 20 iterations. Cells were sampled with replacement to create bootstrap replicates, and clustering was repeated with the same clustering parameters to confirm the same clusters are generated. Heatmap values: ratio of adjusted observation pair counts for each pair of cell subclasses (derived from adjusted Rand index). Each ratio was averaged across bootstrap iterations. High on-diagonal values: cell subclass remains coherent in the bootstrap replicates. High off-diagonal values: corresponding pair of cell subclasses separate in the bootstrap replicates.

**Support File-Figure_5 related to Figure 1-5.**
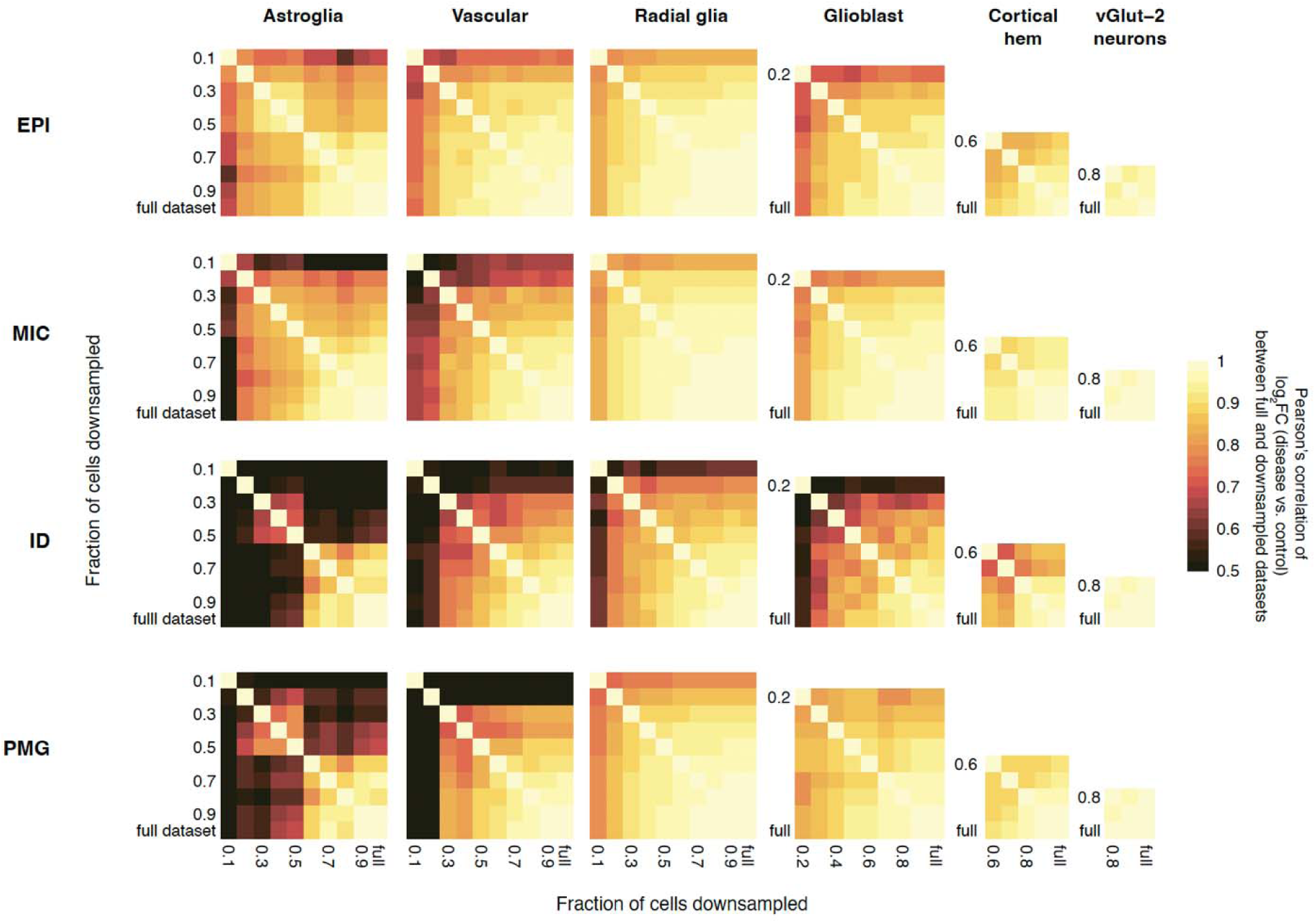
correlations of disease vs. control gene expression fold-changes between full and downsampled datasets. For each sample and each cell subclass, a fraction (0.1 to 0.9 at the step of 0.1) of cells were randomly downsampled, and pseudobulk DE tests with the same setting as in the full dataset using dreamlet were performed. Pearson’s correlations of the log2FC were calculated using all the genes that went into the test for each pair of the downsampled or full datasets.

**Support File-Figure_6 related to Figure 1-5 and Figure S6.**
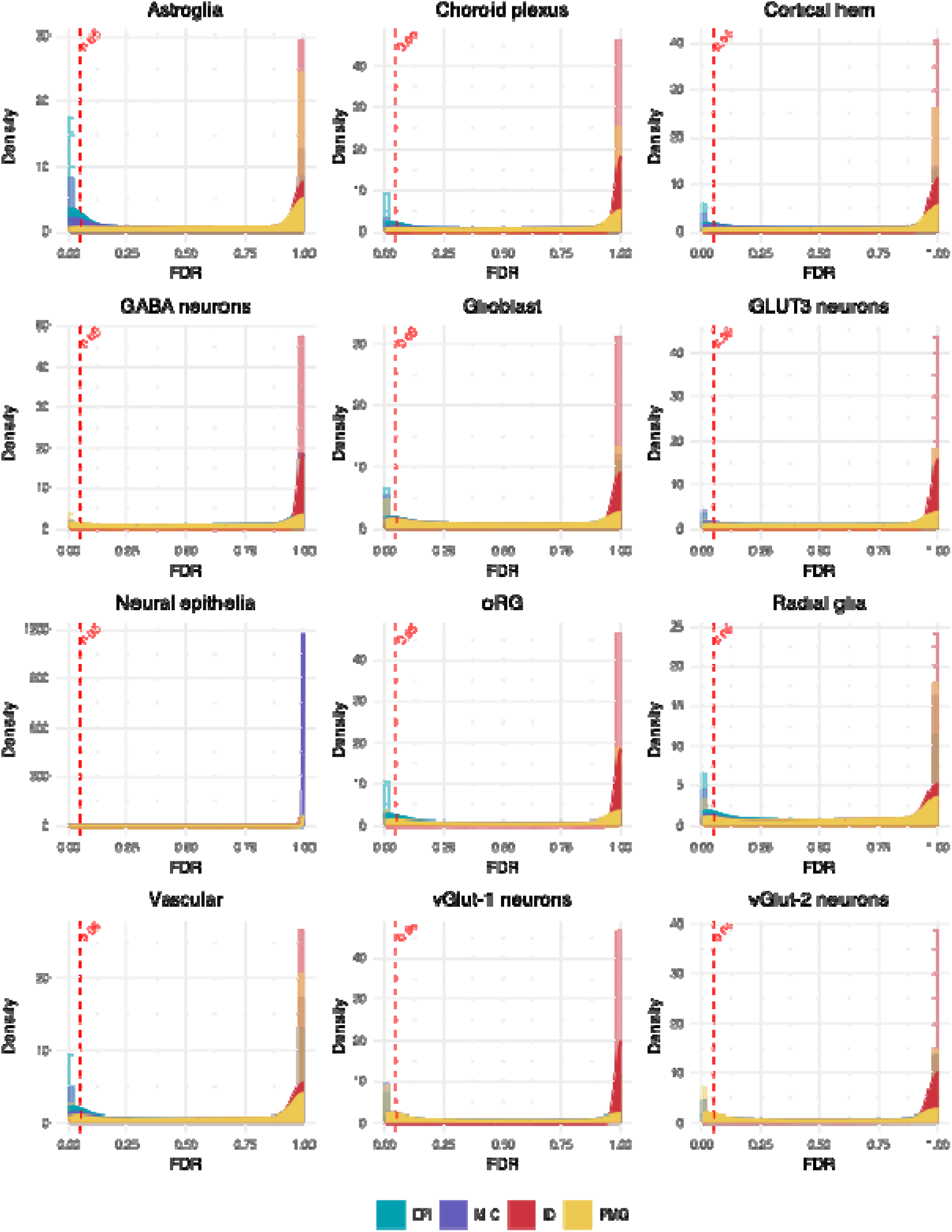
density plots of false discovery rate (FDR) values for NEBULA-LN differential expression results across 16 cell subclasses, grouped by diagnosis. Each panel represents a distinct cell type. Histograms and kernel density estimates are overlaid to visualize the distribution of FDR values per disease. Dashed red lines indicate FDR threshold of 0.05. Across all cell types and conditions, 79,757 gene–cell type– disease combinations passed FDR < 0.05; among these, 15,785 genes are unique.

**Support File-Figure_7 related to Figures 1-5, Figures S5-6.**
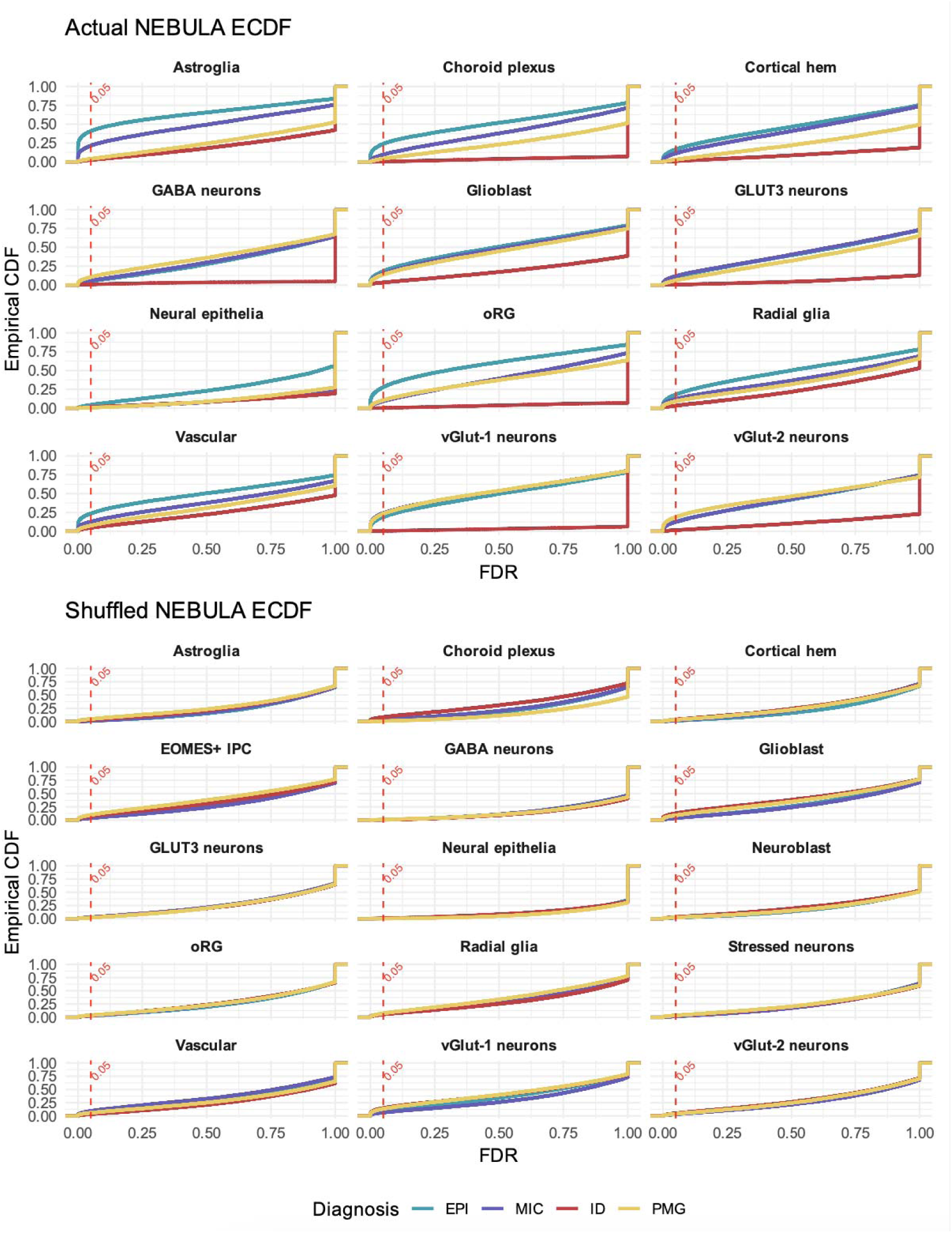
empirical cumulative distribution functions (ECDFs) of NEBULA-LN FDR values for each diagnosis and cell type, comparing disease labels (top) to models fit on randomly shuffled disease labels (bottom). Panels faceted by cell subclass. Curves: ECDF of adjusted p-values (i.e. FDRs) for each diagnosis. Dashed vertical lines: 0.05 significance threshold. Note for example that Astroglia (top-left panel) exhibits a substantially greater proportion of genes with lower FDR for diseases Epilepsy (EPI) and Microcephaly (MIC), and that such patterns are absent in the corresponding shuffled model for Astroglia (bottom-left panel), highlighting differential expression beyond variations explained by cell count differences across disease or cell subclasses.

**Support File-Figure_8 related to Figures S5-6.**
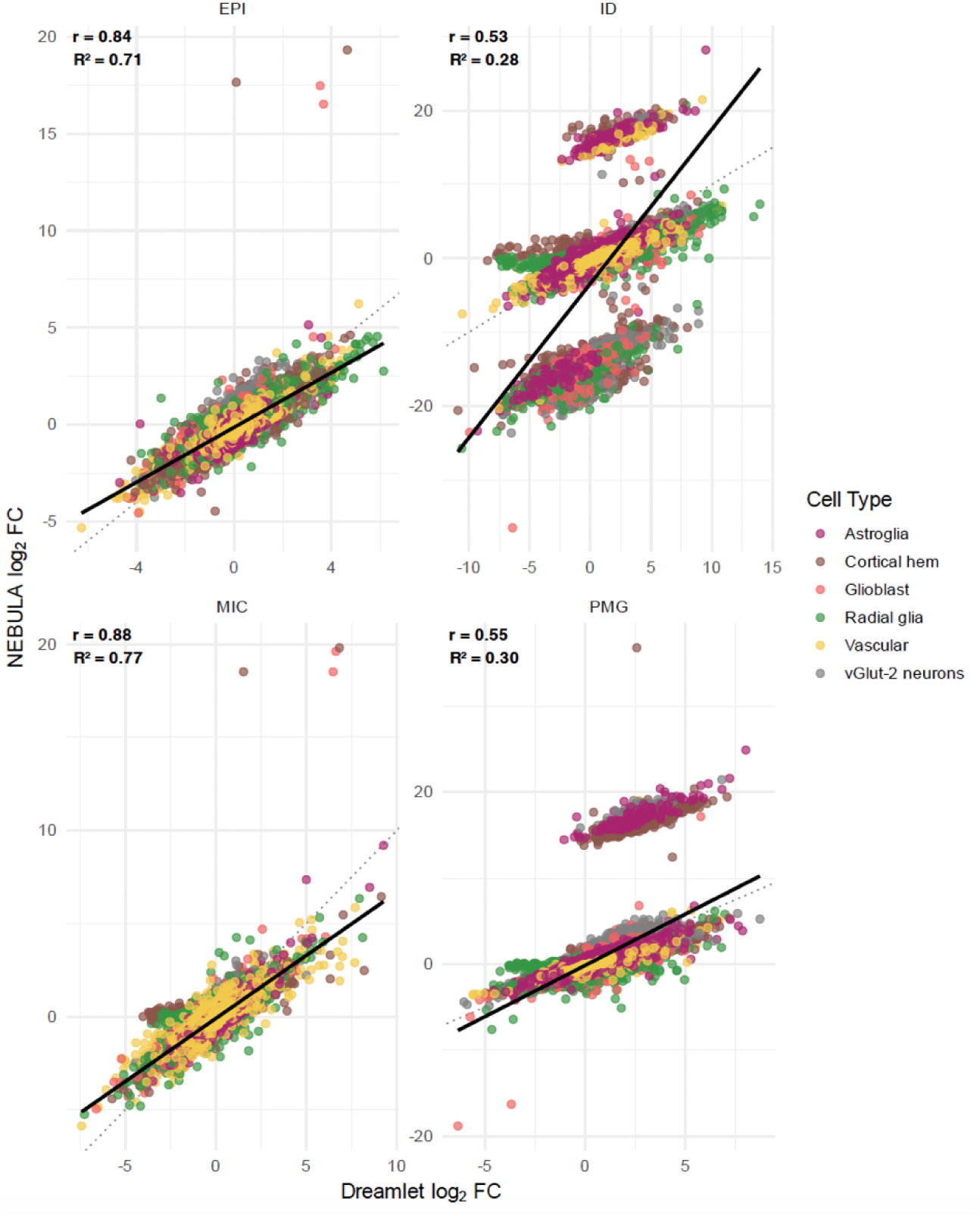
comparison of NEBULA-LN and Dreamlet log_₂_ fold changes (logFC) for differentially expressed genes across four disease conditions (EPI, ID, MIC, PMG). Each panel plots logFC values from NEBULA (y-axis) against Dreamlet (x-axis) for matched gene–cell type combinations. Points are colored by cell subclass. Diagonal dotted lines denote the identity line. Solid lines represent linear fits per disease. Across diseases, NEBULA and Dreamlet logFCs exhibited strong to moderate strong positive correlation, indicating high concordance between single-cell and pseudobulk differential expression analysis.

## Support File_Clinical notes

**For patient brain images**

**NDD-575-4-1 (CW60042)**

Brain MRI images (1-4) findings include white matter atrophy with a thin corpus callosum (CC). Abnormal gyri and cerebral atrophy were observed (images 1-3, axial T1/2-weighted). Medium to mild cerebellar atrophy and thin CC was observed in the sagittal T1-weighted view (image 4). WES available at phs000288.

**NDD-590-4-2 (CW60070)**

Brain MRI images (1-3) findings include severe white matter atrophy, enlarged ventricles, cerebral atrophy and missing CC. Images shown in axial T1/2-weighted views in 1-3 and sagittal views in the lower part of image 1. WES available at phs002032.

**NDD-621-4-2 (CW60188)**

Brain CT images (1-2) findings include slightly enlarged ventricles. All images were collected in the axial view. WES available at phs002032.

**NDD-825-4-2 (CW60430)**

Brain CT images reveal no significant abnormalities.

**NDD-848-4-1 (CW60169)**

Brain MRI images (1-3) findings include severe cerebral atrophy, and reduced volume of cortex. Brain images were collected in axial T2-weighted view (image 1), sagittal T1-weighted view (images 2-3), and coronal T2-weighted view (image 2, lower panel). WES available at phs000288.

**NDD-982-3-4 (CW60142)**

Brain CT and MRI images (1-5) findings include cerebellar atrophy, glio-ependymal cyst, enlarged ventricles, cerebral atrophy, missing CC, and cortical malformations. Images were collected in axial view (images 1-2 and 4-5) and sagittal T1-weighted view (image 3). WES available at phs002032.

**NDD-938-4-7 (CW60054)**

Brain MRI images (1-2) findings include severe cerebral atrophy with a dramatic reduction in the cortical volume. A thin CC and enlarged ventricles were observed. Images were collected in both T1/2-weighted sagittal (image 1) and axial (images 1-2) views. WES available at phs002032.

**NDD-1372-3-1 (CW60408)**

Brain MRI images (1-4) findings include white matter atrophy, enlarged ventricles and cerebral atrophy with reduced cortical volume. Images were collected in both axial T1/2-weighted (images 1-3) and sagittal T1-weighted (image 4) view. WES available at phs003520.

**NDD-1372-3-3 (CW60406)**

Brain MRI images (1-2) findings include moderate level of cerebral atrophy. Images were collected in both axial T1/2-weighted (images 1-2) and sagittal (image 2, left panel) views. WES available at phs003520.

**NDD-1376-4-3 (CW60065)**

Brain MRI images (1-2) findings include mild cerebral atrophy. Images were collected in T1- weighted axial (image 1) and sagittal (image 2) views. WES available at phs000288.

**NDD-1389-5-3 (CW60456)**

Brain MRI and CT images (1-4) findings include white matter atrophy with enlarged ventricles and cerebral atrophy. Images were collected in axial T1/2-weighted (images 1-3) and sagittal T1-weighted (image 2, right panel) views. WES available at phs002032.

**NDD-1426-I-3-4 (CW60056)**

Brain CT images (1-2) findings include slightly enlarged ventricles. All images obtained in axial view. WES available at phs002032.

**NDD-1426-I-3-2 (CW60052)**

Brain MRI images (1-2) findings include cerebral atrophy on both hemispheres. Stacks of axial T1/2-weighted images were collected with a paired sagittal view to reference the location. WES available at phs002032.

**NDD-1599-4-2 (CW60417)**

Brain MRI images findings include severe cerebral atrophy and enlarged ventricles. T1- weighted images were collected in both coronal (upper panel) and sagittal (lower panel) views. WES available at phs002032.

**NDD-1612-6-3 (CW60198)**

Brain CT images (1-2) findings include cerebral atrophy. Images were collected in axial view. WES available at phs003520.

**NDD-1613-4-1 (CW60307)**

Brain MRI images (1-3) findings include atrophies in cerebrum and cerebellum. Both axial T1/2- weighted (images 1 and 3) and sagittal T1-weighted (image 2) views were collected. WES available at phs000288.

**NDD-1654-5-2 (CW60252)**

Brain MRI (1-5 and 7-11) and CT (6) images findings include severe atrophy in cerebellum. MRI Images were collected in axial T1-weighted (images 1, 3, 5, 8 and 10), T2-weighted (images 4, 7, 9, and 11), and sagittal T1-weighted view (image 2). CT images were collected in axial view (image 6). WES available at phs001060.

**NDD-1727-6-3 (CW60505)**

Brain MRI images (1-4) findings include severe cerebral and cerebellar atrophy, missing CC, and enlarged ventricle. Images were collected in axial T1/2-weighted (images 1 and 4), sagittal T1-weighted (image 3), and coronal T2-weighted (image 2) views. WES available at phs001267.

**NDD-1707-III-4-2 (CW60079)**

Brain CT images (1-4) findings suggest cortical malformations. Images were collected in axial (images 1-3) and cornal (image 4) views. WES available at phs002032.

**NDD-1708-6-3 (CW60192)**

Brain CT images reveal no significant abnormalities. Both axial and coronal images were collected. WES available at phs001060.

**NDD-1720-5-2 (CW60193)**

Brain MRI images (1-5) findings include cerebral atrophy, enlarged ventricles, and reduced cortical volume. Images processes for images 1-3 were shown in the lower panel of images. Axial T1-weighted images were shown in image 1; Coronal T2-weighted (left panel) and sagittal T1-weighted (right panel) images were shown in image 2; Axial T1/2-weighted images were shown in image 3; Axial (upper panel) and sagittal (lower panel) T1-weighted images were shown in image 4; Coronal (upper panel) and sagittal (lower panel) T2-weighted images were shown in image 5. WES available at phs001060.

**NDD-1899-5-3 (CW60411)**

Brain MRI and CT images (1-5) findings include mild cerebral atrophy. MRI Images were collected in axial T1/2-weighted (images 1-3), coronal T2-weighted (image 4), and sagittal T1- weighted (image 5) views. CT images were collected in axial view (upper panel, image 2). WES available at phs002032.

**NDD-970-3-1 (CW60058)**

Brain MRI images (1-3) findings include cerebral atrophy. Axial T1/2-weighted (images 1-2) and coronal T2-weighted (image 3) were collected. WES available at phs002032.

**NDD-970-3-2 (CW60084)**

Brain MRI images (1-4) findings include mild cerebral atrophy. Images were collected in axial T1/2-weighted (images 1-2), coronal T2-weighted (image 3), and sagittal T1-weighted (image 4) views. WES available at phs002032.

**NDD-2613-3-4 (CW60305)**

Brain MRI images findings include cerebral atrophy and enlarge ventricles. Images were collected in axial and sagittal T1/2-weighted views. WES available at phs002032.

**NDD-2615-4-2 (CW60280)**

Brain MRI images (1-4) findings include white matter atrophy, cerebral atrophy and reduced cortical volume. Images were collected in axial and coronal T1/2-weighted views. WES available at phs002032.

**NDD-2622-3-18 (CW60621)**

Brain MRI images (1-39) findings include cerebral and cerebellar atrophy. Abnormal gyrifications were observed on both hemispheres. Images were collected in axial, sagittal and coronal T1/2-weighted views. WES available at phs000288.

**NDD-2622-3-20 (CW60623)**

Brain MRI images (1-11) findings include cerebral atrophy, abnormal gyrifications were observed in some of the images with simplified gyrus. Images were collected in axial and sagittal T1/2-weighted views. WES available at phs000288.

**NDD-2629-5-1 (CW60471)**

Brain MRI and CT images (1-8) findings include cerebral atrophy, white matter atrophy, mild enlarged ventricles and reduced cortical volume. MRI Images were collected in axial and sagittal T1/2-weighted views (images 3-8). CT images were collected in axial view (images 1-2). WES available at phs000288.

**NDD-2712-2-1 (CW60510)**

Brain MRI images findings include cerebral atrophy, multiple irregular small gyri, white matter, and cerebellar atrophy, as well as enlarged ventricles. Images were collected in sagittal, axial, and coronal T1/2-weighted views. WES available at phs003520.

**NDD-2745-3-2 (CW60548)**

Brain MRI images (1-2) findings include cerebral atrophy with enlarged ventricles. Images were collected in axial T1-weighted (image 1) and T2-weighted (image 2) views. WES available at phs003520.

**NDD-2920-3-2 (CW60491)**

Brain MRI images (1-3) findings include cerebral atrophy, irregular gyrus, and reduced cortical volume. Images were collected in axial T1/2-weighted (images 2-3) and sagittal T2-weighted (image 1) views. WES available at phs003520.

**NDD-2920-3-3 (CW60492)**

Brain CT images are normal. Images were collected in axial view. WES available at phs003520.

**NDD-2966-3-3 (CW60507)**

Brain MRI images (1-11) findings include cerebral atrophy, irregular gyrus, and cerebellar atrophy. Images were collected in axial, sagittal, and coronal T1/2-weighted views. WES available at phs002621.

**NDD-3074-4-3 (CW60325)**

Brain MRI images (1-5) findings include cerebral atrophy and reduced cortical volume. Axial T1/T2-weighted images were shown in image 1; Coronal T2-weighted (left panel) and axial T2- weighted (right panel) images were shown in image 2; Axial T1-weighted images were shown in image 3; Axial (left panel) and sagittal (right panel) T1-weighted images were shown in image 4; Axial T1/T2-weighted images were shown in image 5. WES available at phs003520.

**NDD-3831-4-1 (CW60261)**

Brain MRI images (1-5) findings include severe cerebral and cerebellar atrophy and enlarged ventricles. Images were collected in axial T1/T2-weighted (images 3-5), coronal T2-weighted (image 1) and sagittal T2-weighted (images 1-2) views. WES available at phs003520.

**NDD-3833-4-2 (CW60651)**

Brain MRI images (1-2) findings include irregular gyrus. Images were collected in axial T1/T2- weighted (images 1-2) and sagittal T1-weighted (image 2, left panel) views. WES available at phs003520.

**NDD-3525-5-4 (CW60626)**

Brain CT findings include slightly enlarged ventricle. Images were collected in axial view. WES available at phs003520.

**NDD-4949-4-2 (CW60327)**

Brain MRI images (1-4) findings include severe cerebral atrophy with reduced cortical volume and enlarged ventricles, irregular gyrus was observed. Images were collected in axial T1/T2- weighted (images 2-4), sagittal T1-weighted (image 1, right panel), and T2-weighted (image 1, left panel) views. WES available at phs003520.

**NDD-4949-4-3 (CW60328)**

Brain MRI and CT images (1-4) findings include severe cerebral atrophy and reduced cortical volume. MRI Images were collected in axial T1/T2-weighted (images 3-4), and sagittal T1- weighted (image 2, lower panel), and coronal T2-weighted (image 2, upper panel) views. CT images were collected in axial view (images 1). WES available at phs003520.

**NDD-4971-4-4 (CW60637)**

Brain MRI images (1-9) findings include severe cerebral atrophy, reduced cortical volume, and enlarged ventricles. Irregular gyrus was observed. Images were collected in axial and sagittal T1/T2-weighted views. WES available at phs001510.

**NDD-5379-3-2 (CW60671)**

Brain CT images findings include cerebral atrophy and enlarged ventricles with a cleft in right hemisphere. Images were collected in axial views. WES available at phs003520.

**NDD-5657-5-1 (CW60669)**

Brain MRI images (1-3) findings include cerebral atrophy and enlarged ventricles. Images were collected in axial T1/T2-weighted (images 1 and 3), coronal T1-weighted (images 2, left panel) and sagittal T1-weighted (images 2, right panel) views. WES available at phs001510.

**NDD-103-3-3 (CW60138)**

Brain MRI images (1-5) findings include cerebral atrophy with enlarged ventricles and reduced cortical volume. Images were collected in axial T1/T2-weighted (images 2-4), sagittal T1- weighted (images 1), and coronal T2-weighted (images 5) views. WES available at phs002032. *Clinical notes corresponding to the images listed in the UCSC website:* https://cells-test.gi.ucsc.edu/?ds=brain-org-ndd

### Epilepsy patients with cortical resection

**EPI-3366**

EPI-3366 is a female individual with epilepsy. Her seizures began at 2 months of age, with a frequency of 20–25 episodes per day despite treatment with multiple medications. Brain MRI revealed findings consistent with focal cortical dysplasia type 2. At the age of 3 years and 3 months, she underwent a lobectomy and corticectomy. Histological analysis confirmed a diagnosis of FCD type 2A. Resected brain tissue was collected for staining.

**EPI-3365**

EPI-3365 is a female individual with epilepsy. Her seizures began at 1 month of age, with a frequency of 10–15 episodes per day despite treatment with multiple medications. Brain MRI revealed findings consistent with focal cortical dysplasia type 2. At the age of 7 months, she underwent a lobectomy. Histological analysis confirmed a diagnosis of focal cortical dysplasia type 2A. Resected brain tissue was collected for staining.

## Support File-Legends for the Supplementary Tables

**Table S1_Data availability for all CIRM-iPSC lines.**

Genome data availability for all CIRM-iPSC lines generated from a cohort of recessive patients based on neurodevelopmental disorders. Patient IDs in the first column, corresponding to the brain images in Support File-1 (https://brain-org-ndd.cells.ucsc.edu/). CIRM IDs listed in second column, corresponding to Fujifilm Cellular Dynamics (https://www.fujifilmcdi.com/cirm-faqs/). Whole Exome/Genome data listed in third column, along with dbGaP (Database of Genotypes and Phenotypes) accession numbers, i.e. “phsxxx.” Total of 318 lines included in this table.

**Table S2_Raw Quantification used in the manuscript.**

**Sheet 1 (Figure 1D–F and Figure S4)-** raw quantifications for panels D to F in Figure 1. NDD- hBOs were stained and quantified for various cellular markers at DIV28 (D and E) and DIV52 (D and F), including SOX2, KI67, Cleaved Caspase-3 (CC3), TBR2, and CTIP2. Marker-positive cells and DAPI-stained nuclei were counted using ImageJ, and the percentage of marker-positive cells was calculated per organoid for each iPSC line. Twelve different hBOs, representing 12 independent biological replicates, were analyzed. The mean of all replicates per category was used to generate the bar plot in panel D. In panels E and F, all replicates are shown as small, faded dots on the PCA plot, with the mean of each iPSC line highlighted as a larger, solid-colored dot. An average of three distinct image regions was quantified per hBO. A total of 47 iPSC lines were assayed, including 13 control lines (e.g., Ctrl1, Ctrl2, Ctrl3, …), 10 microcephaly lines (e.g., MIC1, MIC2, MIC3, …), 8 polymicrogyria lines (e.g., PMG1, PMG2, PMG3, …), 8 epilepsy lines (e.g., EPI1, EPI2, EPI3, …), and 8 intellectual disability lines (e.g., ID1, ID2, ID3, …). S: structural disorders; NS: non-structural disorders; C: control.

- **Sheet 2 (Figure S1E-F)-**Raw quantifications for panel E and F in Supplemental Figure
- Ctrl-hBOs were harvested at DIV28 for immunostaining against SOX2, KI67, and CC3, while CTIP2 and TBR2 were stained at DIV52. The number of marker-positive and DAPI-positive cells was counted at each time point (DIV28 or DIV52), and the percentage of marker-positive cells was calculated. Twelve different hBOs, representing 12 independent biological replicates, were used to generate the bar plot in panel E. The box plot in panel N shows the percentage of each cellular marker across replicates. Sample variation was calculated and plotted in panel S2.
- **Sheet 3 (Figure 2B-E)-**Raw quantifications for panel B to E in Figure 2. MIC-hBOs were harvested and stained at DIV28 (orange) and DIV52 (blue) for SOX2, KI67, and CC3, while CTIP2 and TBR2 were harvested and stained at DIV52. The number of marker-positive and DAPI-positive cells at DIV28 or DIV52 was counted, and the percentage of marker-positive cells was calculated. Twelve different hBOs, representing 12 independent biological replicates, were used for the analysis.
- **Sheet 4 (Figure S5B)-**Raw quantifications for the size of hBOs across disease categories. The diameters of the hBOs were measured at DIV28 (orange), DIV52 (blue), and DIV80 (pink) for each of category. Twelve independent organoids were measured for each individual iPSC line (from -R1 to -R12), and the mean and standard deviation (SD) were calculated for statistical analysis and plotting. A total of ten control lines (Ctrl1 to Ctrl10, green), nine microcephaly lines (MIC1 to MIC9, purple), eight polymicrogyria lines (PMG1 to PMG8, yellow), eight epilepsy lines (EPI1 to EPI8, cyan), and six intellectual disability lines (ID1 to ID6, red) were assayed.
- **Sheet 5 (Figure 3B-E)-**Raw quantifications for panel B to E in Figure 3. PMG-hBOs were harvested and stained at DIV28 (orange) and DIV52 (blue) for SOX2, KI67, and CC3. CTIP2 and TBR2 were harvested and stained at DIV52. The number of marker-positive and DAPI-positive cells were counted, and the percentage of positive cells was calculated. Twelve different hBOs, representing 12 independent biological replicates, were analyzed.
- **Sheet 6 (Figure 4B-E)-**Raw quantifications for panel B to E in Figure 4. EPI-hBOs were harvested and stained at DIV28 (orange) and DIV52 (blue) for SOX2, KI67, and CC3. CTIP2 and TBR2 were harvested and stained at DIV52. The number of marker-positive and DAPI-positive cells was counted, and the percentage of positive cells was calculated. Twelve different hBOs, representing 12 independent biological replicates, were used.
- **Sheet 7(Figure S6B and G)-**Raw quantifications for panel B and G in Figure S6. EPI- hBOs and Ctrl-hBOs were harvested and stained at DIV65 for GFAP, an astrocyte marker. The number of astrocytes was quantified by measuring the GFAP+ area and manually counting based on astrocytic morphology. DAPI-positive cells were counted to identify the cells. Data were tested for normality prior to analysis to ensure the correct statistical tests were used.
- **Sheet 8 (Figure 5B-E)-**Raw quantifications for panel B to E in Figure 5. ID-hBOs were harvested and stained at DIV28 (orange) and DIV52 (blue) for SOX2, KI67, and CC3. CTIP2 and TBR2 were harvested and stained at DIV52. The number of marker-positive and DAPI-positive cells was counted, and the percentage of positive cells was calculated. Twelve different hBOs, representing 12 independent biological replicates, were used.
- **Sheet 9 (Figure S7B-E)-**Raw quantifications for panel B to E in Figure S7. Additional ID- hBOs were harvested and stained at DIV28 (orange) and DIV52 (blue) for SOX2, KI67, and CC3. CTIP2 and TBR2 were harvested and stained at DIV52. The number of marker-positive and DAPI-positive cells was counted, and the percentage of positive cells was calculated. Six different hBOs, representing six independent biological replicates, were used.
- **Sheet 10 (Figure S9E)-**Raw quantifications for panel E in Figure S9. *NARS1* healthy, mutant, and corrected hBOs were harvested at DIV28 (orange) to measure their size. The diameters of the hBOs were measured for each genotype. Three different hBO generations, representing three independent biological replicates, were used.
- **Sheet 11 (Figure S10B)-**Raw quantifications for panel C in Figure S10. *ACTL6B* healthy, mutant, and corrected hBOs were harvested at DIV28 for qPCR evaluation of the early response genes *FOS*, *DUSP5*, *JUN*, *VGF*, *FOSB*, and *FOSL2.* Relative mRNA expression was calculated and plotted, with *GAPDH* used as the reference. Three independent hBOs were measured.
- **Sheet 12 (Support File-Figure 2B)-**Raw quantifications for panel B in Support File-Figure 1B. NDD-hBOs were harvested and stained for TBR1 at DIV80. The number of TBR1+ and DAPI+ cells was counted. The percentage of TBR1+ cells was calculated and plotted. Three independent hBOs were measured.

**Table S3_scRNAseq metadata.**

Metadata for the scRNA-seq related to Figures 1-5 and S2 to S6: A total of 155,324 cells are listed in the table. The columns from left to right are as follows:

- **A**_cell_id: 10X cell barcode for each cell.
- **B**_sample: Each sample for the single-cell RNA-seq.
- **C**_num_UMI_raw: Raw number of UMIs for each cell obtained from Cell Ranger.
- **D**_num_genes_raw: Raw number of genes for each cell after processing by Cell Ranger.
- **E**_diagnosis: Disease categories for each sample.
- **F**_sex: Sex for each of the samples and cells assayed.
- **G**_batch: Batch for the single-cell RNA-seq experiment corresponding to different samples.
- **H**_percent_mitochondrial_reads: The percentage of mitochondrial reads for each cell.
- **I**_doublet_scores: Scores indicating the presence of two cells in one droplet in the single-cell RNA-seq experiment.
- **J**_num_UMI_SCtransformed: The number of normalized UMIs in each cell. Normalization was performed using the ‘SCtransformed’ function in Seurat.
- **K**_num_genes_SCtransformed: The number of genes after normalization in Seurat.
- **L**_Seurat_cluster: The cluster assignment for each cell using the Seurat pipeline.
- **M**_annotated_region: The predicted region for each cell based on comparison with the human fetal brain dataset.
- **N**_annotated_class: The annotated class for each cell based on transcriptional features.
- **O**_annotated_subclass: The annotated subclass for each cell based on transcriptional features.
- **P**_annotated_cell_type: The annotated cell type for each cell.
- **Q-R**_average_predicted_age/age_bracket: The average predicted age and corresponding age bracket for each cell, predicted by the human fetal brain dataset.
- **S-T**_monocle3_partition/pseudotime: The fate trajectory for each cell predicted using Monocle3. Partition and pseudotime were calculated for plotting.

**Table S4_scRNAseq cluster annotation**

Description of the annotation of different cell clusters, related to **Figures 1-5 and S2-3**: A total of 59 clusters were annotated based on highly expressed regional markers for different brain regions/sub-regions (**Columns B-D**), and highly expressed cell type markers for different cell classes, subclasses, and cell types (**Columns E-H**). The status of cellular cycling was predicted and indicated as True/False (**Column I**). The age/age bracket for each cluster was calculated (**Columns J-K**).

**Table S5_scRNAseq marker genes for each cluster.**

The expression of marker genes for each cluster was calculated and statistically analyzed, related to **Figures 1-5**. Clusters and genes are listed in **Columns A** and **B**. Log2 fold change of the average expression for each gene was calculated (**Column C**). The percentage of cells where the gene is detected in the tested cluster (**Column D**) and in other clusters (**Column E**) was calculated. p-values (**Column F**) and Bonferroni-adjusted p-values (**Column G**) were calculated to assess significance.

**Table S6_Classifier model performance for LDA**

**Table S7_Statistics in Figures 2-5**.

Quantification calculations and statistics for the immunostaining in Figures 2-5.

- **Sheet_1: Quantifications.** Disease category, sample, timepoint, and marker assayed are listed in **columns A** to **D**. Proportion mean and proportion median are calculated in **columns E** and **F**. Proportion standard deviation (SD) and proportion standard error of the mean (SEM) are calculated in **columns G** to **H**. Proportion confidence interval (CI) is calculated in **column I**.
- **Sheet_2: Statistics.** Statistics corresponding to the quantification in Sheet_1 are as follows: Marker genes are listed in **column A**; two different time points are listed in **column B**; Proportion_mean_DIVX_Marker is listed in **column C**. A two-sided t-test was used for the statistical analysis (**columns D and E**). Group 1 results, from the control group, are listed in **column F**, and group 2 results, from disease categories, are listed in **column G**. The total number of samples is listed as n1 and n2 in **columns H** and **I**, respectively. Calculations for statistics are shown in **columns J** to **Q**.

**Table S8_scRNAseq cell class proportion test by propeller.**

Raw proportion propeller test results for scRNA-seq cell classes, related to Figures 2 to 5:

- **Sheet 1:** The number of cells in each cell class across different samples.
- **Sheet 2:** The proportion of cells in each cell class across different samples.
- **Sheet 3:** The logit-transformed proportion for each cell class across different samples.
- **Sheet 4:** The asin-transformed proportion for each cell class across different samples.
- **Sheet 5:** Statistical test using the logit-transformed proportion in Ctrl vs. MIC.
- **Sheet 6:** Statistical test using the asin-transformed proportion in Ctrl vs. MIC.
- **Sheet 7:** Statistical test using the logit-transformed proportion in Ctrl vs. PMG.
- **Sheet 8:** Statistical test using the asin-transformed proportion in Ctrl vs. PMG.
- **Sheet 9:** Statistical test using the logit-transformed proportion in Ctrl vs. EPI.
- **Sheet 10:** Statistical test using the asin-transformed proportion in Ctrl vs. EPI.
- **Sheet 11:** Statistical test using the logit-transformed proportion in Ctrl vs. ID.
- **Sheet 12:** Statistical test using the asin-transformed proportion in Ctrl vs. ID.

**Table S9_scRNAseq Dreamlet pseudo-bulk differential expression (DE) genes.**

List of genes that are differentially expressed in different cell classes across samples: (**Sheet 1:** MIC vs. Ctrl; **Sheet 2:** PMG vs. Ctrl; **Sheet 3:** EPI vs. Ctrl; **Sheet 4:** ID vs. Ctrl).

**Table S10_ Differentially Expressed Genes with NEBULA**

List of genes that are differentially expressed: (**Sheet 1:** full list; **Sheet 2:** FDR < 0.05; **Sheet 3:** FDR < 0.2;).

**Table S11_Selected genes in cell subclass pseudobulk wilcox test raw pvalues add fdr**

**Table S12_Primers used in this study.**

## Support File-Methods for scRNA-seq analysis in details

### scRNA-seq data preprocessing, clustering and cell type annotation

scRNA-seq reads were mapped to the human GRCh38 genome with pre-built gene annotations (refdata-gex-GRCh38-2020-A) using 10x Genomics Cell Ranger (v7.1.0) with the default parameters (where the “--include-introns” flag was turned on to include intronic reads towards UMI counting). The raw feature-by-droplet matrices generated from the “cellranger count” module were then fed to CellBender (v0.2.0)2 to distinguish cell-containing from cell-free droplets and retrieve noise-free cell-by-gene quantification tables using the following parameters: “--expected-cells 1000 --total-droplets-included 25000 --model full --epochs 150 --cuda --low-count-threshold 15 --fpr 0.01 --learning-rate 0.0001 --posterior-batch-size 5 --cells-posterior-reg- calc 50”. Samples that retained no cells after CellBender were excluded from the downstream analysis. Scrublet (v0.2.3)3 was then applied to the CellBender-generated h5 file for each sample to compute the doublet score for each cell with an expected doublet rate of 0.06. The doublet score cutoff was set to 0.2 based on the simulated doublet histogram of doublet scores, and this threshold worked well for all samples.

Next, Seurat (v4.3.0) was used to create a merged object containing all the samples, and to perform the following downstream processing. Cells with fewer than 500 detected genes, more than 5% reads mapped to the mitochondrial genome, or a doublet score greater than 0.2 were excluded from further analysis. Sctransform v2 normalization5 was performed on the remaining cell, with the mitochondrial mapping percentage and the sequencing batches set as the confounding sources of variation to be removed. Principal component analysis (PCA) was run with the top 3000 highly variable genes in the normalized cell-by-gene matrix, and the first 50 principal components were used to run Harmony (v0.1.1) integration with sample IDs and sequencing batches set as covariants. Then, a k-nearest-neighbor graph was constructed with k set as 30, and the cells were clustered using Leiden clustering with a resolution of 1.

To remove putative stressed cells in brain organoid samples due to insufficient oxygen transport, we applied granular functional filtering, Gruffi7, to the clustered scRNA-seq dataset mentioned in the previous step. Gruffi first partitioned cells into groups of 100-200 cells by finding the right granule resolution, then computed Gene Ontology (GO) pathway scores per cell, and took the average GO scores per granule (normalized by respective cell numbers). The GO terms used for stressed cell filtering were glycolytic process (GO:0006096), response to endoplasmic reticulum stress (GO:0034976), and gliogenesis (GO:0042063). We then used the automatic stress threshold estimation by Gruffi to filter out cells with high glycolysis (estimated GO score cutoff: 2.807) and high ER stress (estimated GO score cutoff: 1.731) but also keep cells with high gliogenesis (estimated GO score cutoff: 1.953). This process removed a total of 21,278 cells, which took up 1.36-34.9% of cells from each sample. Then, the aforementioned Sctransform v2 normalization, PCA, Harmony integration, and Leiden clustering steps were run on the remaining cells, using the same set of parameters except that the Leiden clustering resolution was set to 2. This gave rise to 59 clusters of 155,323 cells, and UMAP (Uniform Manifold Approximation and Projection) was used for visualization.

Cell type annotation was performed in a semi-automatic way. First, marker genes for each cluster were identified using Seurat’s “FindAllMarkers” function with the following parameters: only.pos = TRUE, test.use = "wilcox", min.pct = 0.25, logfc.threshold = 0.25. Next, we transferred the annotated labels from a recently published cell atlas of the first-trimester developing human brain to our dataset. To avoid batch effects arising from different single-cell sequencing technologies, we only used samples prepared by 10X v3 chemistry in the reference dataset. We also removed cell clusters that were annotated with non-central nervous system cell classes: erythrocyte, fibroblast, immune, neural crest, and placodes. The reference dataset was randomly subsetted to have a maximum of 500 cells per original cluster to improve computational efficiency. This retained 251,439 cells in the reference dataset. The cell-by-gene count table from the reference dataset was imported as a Seurat object and normalized using the Sctransform v2 normalization method. Then, we used Seurat’s “FindTransferAnchors” and “TransferData” functions to transfer the annotated labels from the reference (cluster IDs, brain regions, brain subdivisions, cell classes, and developmental ages) to our dataset. For each of 59 clusters defined in our dataset and each of 5 label categories, we counted the number of cells by the top predicted label and kept the top 3 labels for further consideration. The final annotations were then jointly compared using these labels with the marker genes found in the first step and known regional and cell-type markers. The predicted age for each cluster was determined by the average predicted age across the cells within the cluster (5-8, Early; 8-10, Mid; >10, Late). scRNA-seq data were mapped to a human cortical developmental reference atlas based on a harmonized meta-analysis from published single-cell datasets, integrating fetal and early postnatal human brain samples described in Nano et al. 2025, which captures a high-resolution trajectory of cortical development across gestational and early postnatal stages. Seurat (v4.3) label transfer framework was used to project hBO single-cell transcriptomes onto this reference. The developmental meta-atlas (clean.Dev_metaatlas_integrated.rds) and pre- processed annotated hBO dataset (NDD_scRNA_postQC_scTransformed_v2_remove_stressed_cells_annotated.rds) were harmonized by ensuring the same feature space using SelectIntegrationFeatures. The top 2,000 most variable genes shared in both datasets were selected. Dimensionality reduction was performed on the reference using PCA (dims = 1:30), followed by UMAP embedding with return.model = TRUE to enable subsequent query mapping. Anchors between the reference and query were identified using FindTransferAnchors with the PCA space of the reference (reference.reduction = "pca") and transferred developmental labels (gestational week and developmental period) to the query using MapQuery. The reference dataset has a total 599,211 cells with developmental labels. All cells in query dataset obtained the transferred label with a total of 155,323 cells. The mapping returned predicted labels for Gestational Week (GW) and Developmental Period (Dev_Period_Kang2011) for each cell in NDD-hBOs scRNA-seq dataset, allowing us to classify them into eight distinct developmental windows: GW 6–10, GW 10–12, GW 12–15, GW 15–18, GW 18–21, GW 21–26, GW 26–40, and 8 months postnatal as shown in Figure 1C. Major cell types were then annotated for each of the development window as shown in Figure S3A.

NDD_scRNA_postQC_scTransformed_v2_remove_stressed_cells_annotated.rds was subset based on diagnosis labels. Cells were classified into experimental groups according to the pathDx metadata field, annotated as “Healthy”, “Epilepsy”, “Microcephaly”, “Polymicrogyria”, and “Intellectual disability”. Four independent Seurat objects were generated using Seurat (v4.3) for comparison. There are: NDD_C_E (Healthy vs. Epilepsy), NDD_C_M (Healthy vs. Microcephaly), NDD_C_P (Healthy vs. Polymicrogyria), and NDD_C_ID (Healthy vs. Intellectual Disability). Each subset was reprocessed with SCTransform normalization, dimensional reduction, UMAP embedding, and clustering with a resolution of 0.8. This results in 33 clusters of 63937 cells in NDD_C_E, 29 clusters of 64134 cells in NDD_C_M, 33 clusters of 89777 cells in NDD_C_P, and 28 clusters of 73384 cells in NDD_C_ID. Merged UMAP was plotted for NDD_C_E, makers genes evident astrocytic states were plotted as shown in Figure S7D-E.

### Testing for cell type proportion differences across diagnosis

To test whether the cell type proportions are different between organoids originating from disease and control subjects, we applied Propeller (implemented in the R package speckle, v0.99.7), a robust and flexible method that leverages biological replication to find statistically significant differences in cell type proportions between groups10. Cell type proportions were first logit transformed or arcsine transformed, and the proportion differences between each disease and control were tested using the “propeller.ttest” function, with sex added as a covariate in the linear model. Significant proportion differences were called with groups with FDR < 0.2 in either the logit transformed test, or the arcsine transformed test.

### Differential expression test of pseudobulk gene expression in scRNA-seq

We first converted the Seurat object to a SingleCellExperiment object, then applied Dreamlet (v0.99.6)11 to perform differential expression (DE) tests on the pseudobulk count matrix generated from each cell subclass. Genes that were not expressed in any of the chosen cells were removed. Pseudobulk counts were aggregated by cell subclass and by sample using the raw counts from the “RNA” slot in the original Seurat object. DLX2+ IPC, mixed neurons and muscle cells were removed from the following analysis due to insufficient number of cells per sample (at least 20 cells per sample in at least 10 samples across diagnosis groups). Pseudobulk counts were then normalized and applied voom with Dream weights using the “processAssays” function with the following parameters: min.cells = 5, min.count = 5, min.samples = 4, min.prop = 0.3. Next, DE tests for each disease vs. control were performed using a linear model with sex and batch added as covariates. The full list of DE results are provided in Table S9, Downsampled DE as Zip file. Gene Set Enrichment Analysis (GSEA)12,13 was performed with clusterProfiler (v4.4.4)14,15 using the log2FC from all the genes included in the Dreamlet test for each cell subclass and each disease vs. control group. Gene sets tested in GSEA were Gene Ontology Biological Processes terms with more than 10 genes and less than 250 genes.

